# Small-hairpin RNAs cause target-independent microRNA dysregulation in neurons and elicit global transcriptomic changes

**DOI:** 10.1101/2020.07.30.229443

**Authors:** Rafi Kohen, Katherine T. Baldwin, Patricia M. Garay, Takao Tsukahara, Alex Chen, Corey G. Flynn, Craig Johnson, Xiao-Feng Zhao, Michael A. Sutton, Shigeki Iwase, Roman J. Giger

## Abstract

Small hairpin RNAs (shRNAs) allow highly efficient gene knockdown. Here we employed different shRNAs to knock down the reticulon RTN4A/NogoA in primary neurons. Depletion of NogoA correlates with altered synaptic protein composition and spontaneous neurotransmission. However, similar phenotypes are not observed upon genetic deletion of *Nogo* or its receptors. Step-wise introduction of mismatches in the seed region of shNogoA provides further evidence that synaptic phenotypes are NogoA-independent. RNA sequencing revealed global changes in the neuronal transcriptome of cultures transduced with the original shNogoA or closely related variants. Transcriptomic changes are shRNA seed sequence dependent, but not target-specific. Parallel sequencing of small non-coding RNAs revealed dysregulation of microRNAs. Computational analysis shows that the altered miRNA composition correlates with changes in mRNA expression and preferentially affects protein-protein networks that function at synapses. Thus, off-target effects associated with shRNAs are an inherent property, and in particular, altered miRNA composition needs careful consideration.

## INTRODUCTION

RNA interference (RNAi) is a powerful and commonly used research tool, broadly employed for gene silencing and loss-of-function studies. Strengths of this technique are the simple design, easy production of double-stranded RNA molecules, and broad applicability *in vitro* and *in vivo* (Hannon, 2002; McCaffrey et al., 2002; Mello and Conte, 2004). In recent years, RNAi has been used to devise therapeutics for various disorders (Howard, 2009; Pecot et al., 2011; Singh et al., 2018).

The molecular basis of RNAi-mediated gene silencing has been studied extensively. The protein components involved in RNAi-mediated post-transcriptional gene regulation are broadly shared with microRNA (miRNA)-mediated gene regulation (Bernstein et al., 2001; Han et al., 2004; Lam et al., 2015; Lee et al., 2003; Matsuyama and Suzuki, 2019; Moore et al., 2010; Yi et al., 2003). miRNAs are small non-coding RNAs that influence messenger RNA (mRNA) stability and protein synthesis through interaction with the 3’-untranslated region (UTR) of mRNAs (Bartel, 2009; He and Hannon, 2004; Song, 2020). A large number of miRNAs have been identified that regulate neuronal protein synthesis, and some miRNAs are preferentially localized to specific subcellular compartments, including axons, dendrites, and synapses (Kye et al., 2007; Schratt, 2009). At synapses, local protein synthesis controls synaptic morphology and function. Some miRNAs are processed in a neuronal activity-dependent manner and critical for brain health (Cohen et al., 2011; Hu and Li, 2017; Rajgor et al., 2020; Sambandan et al., 2017; Schratt, 2009; van Spronsen et al., 2013).

Experimentally, RNAi can be delivered as small interfering RNA (siRNA) duplexes or as vector-based short hairpin RNA (shRNA) that contains a loop structure, which, upon transcription, is processed into a double-stranded siRNA. An advantage of shRNA is vector-based expression, permitting use of cell-type specific or inducible promoters. When combined with viral vector-mediated gene transfer, this allows efficient and long-lasting gene knockdown (Brummelkamp et al., 2002; Krichevsky and Kosik, 2002; Nielsen et al., 2009; Tönges et al., 2006). RNAi-mediated gene silencing is a multi-step process that results in the degradation of the target mRNA. While siRNA duplexes are localized to the cytoplasm, shRNAs are synthesized in the nucleus, where they form a hairpin structure. The initial transcript is further processed by Drosha and its double-stranded RNA-binding partner Dgcr8 to generate a pre-shRNA, which is then translocated into the cytoplasm and subsequently cleaved by Dicer into a double-stranded siRNA. The siRNA is subsequently loaded onto the RNA-induced silencing complex (RISC). The guide strand of the activated RISC–siRNA complex binds via Watson-Crick base-pairing to the target mRNA, which leads to RNA degradation and gene silencing (Han et al., 2004; He and Hannon, 2004; Lee et al., 2003; Matsuyama and Suzuki, 2019; Yi et al., 2003). A limitation of this technique is that partial sequence complementarity of the siRNA sense or antisense strand to non-target RNAs leads to off-target effects. Thus, extensive validation of siRNA-mediated gene knockdown is a key requirement (Alvarez et al., 2006; Baek et al., 2014; Bartoszewski and Sikorski, 2019; Jackson et al., 2003).

Here, we used lentiviral (LV) vector-mediated delivery of shRNA constructs into primary forebrain neurons to study the function of the reticulon family member RTN4a/NogoA. NogoA is a potent inhibitor of neurite outgrowth *in vitro* and restricts structural neuronal plasticity in the healthy and injured mammalian CNS *in vivo* (McGee et al., 2005; Mironova and Giger, 2013; Schwab and Strittmatter, 2014; Zemmar et al., 2014a). Additional work has found that the Nogo receptor 1 (NgR1) is a negative regulator of activity-dependent synaptic transmission (Lee et al., 2008; Raiker et al., 2010b; Wills et al., 2012). NogoA harbors growth inhibitory motifs called Nogo-Δ20 and Nogo-66, both of which are sufficient to restrict structural and functional neuronal plasticity (Delekate et al., 2011; Kempf and Schwab, 2013; Oertle et al., 2003; Pernet and Schwab, 2012; Pradhan et al., 2010; Raiker et al., 2010a; Zagrebelsky et al., 2010; Zemmar et al., 2017; Zemmar et al., 2014b). These findings indicate that NogoA may function as an important coordinator of structural and functional neuronal plasticity. Whether NogoA plays a role in spontaneous synaptic activity or bidirectional homeostatic synaptic scaling of neuronal networks has not yet been examined. To address these questions, we employed synthetic shRNA constructs for acute knockdown of *NogoA* in developing neurons and found striking changes in synaptic protein composition and function. However, these changes were not replicated following genetic deletion of *Nogo* or its high-affinity receptors. To explore the underlying mechanisms, we employed a combination of biochemical, genetic, and functional genomics studies. We find that neuronal shRNA expression results in global transcriptomic changes that are independent of target gene expression, but shRNA seed sequence dependent. We constructed closely related shRNA vectors bearing mismatches in the seed sequence and found that they all disrupt miRNA composition, including numerous miRNAs that regulate synaptogenesis and neurotransmission. Dysregulation of miRNAs may be an inherent property of shRNAs that needs careful consideration.

## RESULTS

### Synaptic defects following shRNA-mediated knockdown, but not following genetic deletion, of NogoA

For acute NogoA loss-of-function studies in rat hippocampal cultures, we tested four small hairpin RNA (shRNA) constructs, including three commercially available shRNAs and a previously characterized construct (Zagrebelsky et al., 2010) **(Table 1)**. Lentiviral (LV) vectors were used for neuronal transduction. Expression of shRNAs was driven by the H1 or U6 promoter, two weak RNA polymerase III promoters that yield low level shRNA expression to reduce potential off-target effects (Ma et al., 2014). As controls, empty LV with a pLentiLox3.7-GFP backbone (LV-GFP) and a scrambled shRNA (LV-shScram) were used. Primary hippocampal neurons were transduced with LV vectors on day *in vitro* (DIV) 1-3, prior to synapse formation. On DIV 14-17, cultures were lysed and subjected to Western Blot (WB) analysis. Three of the four shRNAs targeting NogoA (LV-shNogoA#1, LV-shNogoA#3, and LV-shNogoA#4) were highly efficient, and showed greater than 90% knockdown of NogoA at the protein level. In contrast, cultures transduced with LV-shNogoA#2, LV-GFP, and LV-shScram did not show any changes in NogoA protein expression (**Figure 1A**). Further analysis revealed that NogoA knockdown correlated with a significant reduction in the glutamatergic AMPA receptor subunit GluA1 and phospho-S6K (pS6K). Protein levels of the NogoB isoform, encoded by a transcript not targeted by the shNogoA, remained unchanged (**Figures 1A and S1A**). For subsequent mechanistic studies, we decided to use LV-shNogoA#1, a previously characterized construct (Zagrebelsky et al., 2010). The kinetics of shRNA-mediated NogoA knockdown and GluA1 decrease were further investigated in hippocampal cultures transduced on DIV3, DIV10, and DIV14. Cultures were lysed on DIV17 and analyzed by WB. NogoA was significantly reduced when cultures were transduced on DIV3 and DIV10, but not on DIV14. Furthermore, loss of NogoA consistently coincided with a significant reduction in GluA1 (**Figure S1B**).

**Figure 1.**
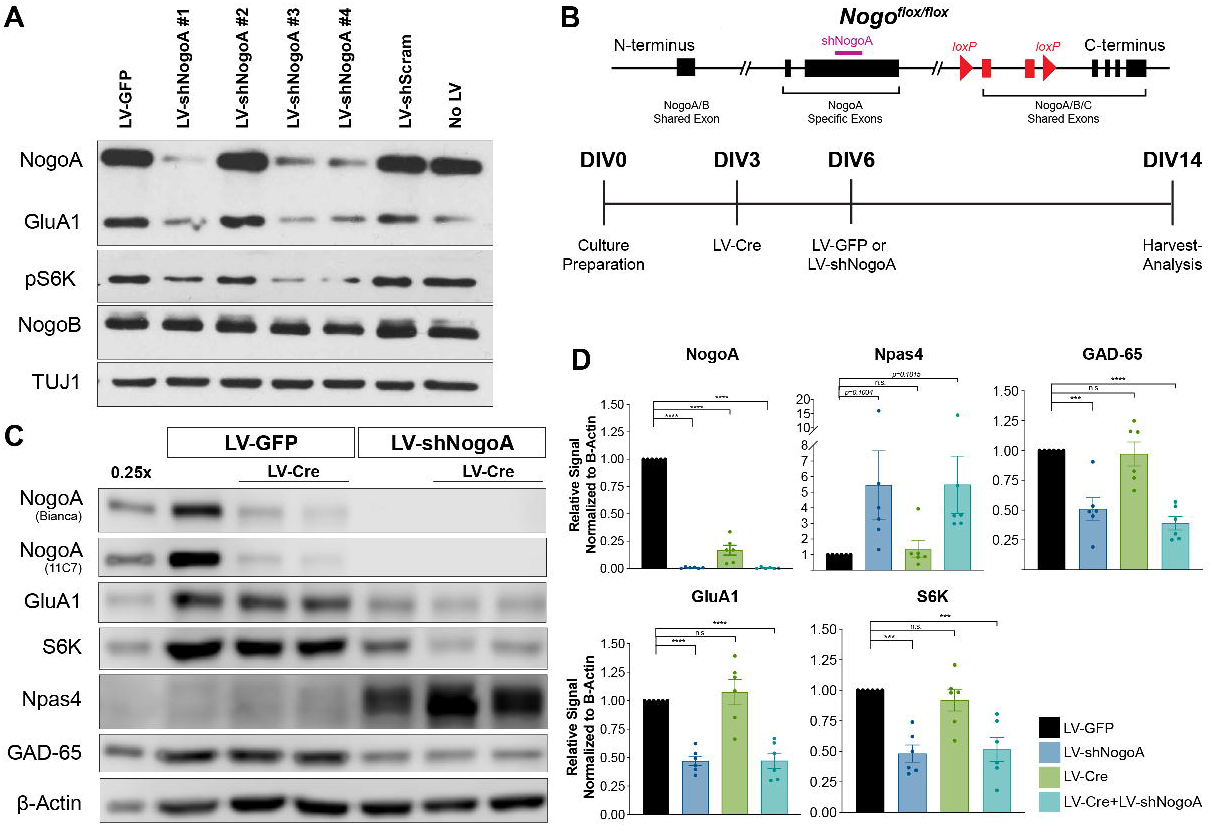
**(A)** Western blot analysis of E18 rat hippocampal neurons. Cultures were transduced on DIV10 and harvested on DIV17. Lentiviral vectors (LV) included LV-GFP, LV-shNogoA#1, LV-shNogoA#2, LV-shNogoA#3, LV-shNogoA#4, LV-shScram. NogoA was probed with the commercial (R&D) antibody. **(B)** Diagram of the *Nogo^flox/flox^* conditional locus, where exons flanked by *loxP* sites (red) or targeted by the shNogoA (violet) are indicated. Timeline is shown for experiments in C-D. **(C)** Western blot analysis of *Nogo^flox/flox^* hippocampal cultures, first transduced with LV-Cre and then with LV-GFP or LV-shNogoA. The 0.25x sample loading is shown to ensure analysis within the linear range of detection. NogoA was independently probed with a monoclonal anti-NogoA (11C7) and a polyclonal anti-NogoAB (Bianca) antibody. **(D)** Quantification of proteins detected by Western blotting. Transduction groups are labeled according to the color key. Signal intensity was first normalized to the β-Actin, then to LV-GFP. Data are presented as mean ± SEM from *n* = 6 biological replicates. \*\*\**p_adj_* < 0.001; \*\*\**p_adj_* < 0.0001, by ordinary one-way ANOVA followed by Dunnett’s multiple comparisons test. n.s., not significant.

Next, we attempted a rescue experiment with an shNogoA-resistant epitope tagged human transgene (LV-hNogoA-Myc). Upon transduction of HEK293T cells with LV-hNogoA-Myc, recombinant NogoA was strongly expressed; however, despite multiple attempts, we failed to express recombinant NogoA in primary neurons at levels comparable to endogenous protein (data not shown). As an independent and more stringent control, we obtained mice carrying a conditional allele for *RTN4/Nogo (Nogo^flox/flox^),* and prepared primary hippocampal cultures (**Figure 1B**) (Meves et al., 2018). DIV3 neurons were transduced with LV-Cre, and cell lysates analyzed by WB on DIV14. In *Nogo^flox/flox^* cultures transduced with LV-Cre, both NogoA and NogoB isoforms were reduced by ≥80% (**Figures 1C-D** and **S1C-D**); however, GluA1 and total S6K levels were similar to cultures transduced with LV-GFP (**Figure 1C-D**).In a parallel experiment, *Nogo^flox/flox^* cultures were transduced first with LV-Cre on DIV3 and three days later with either LV-GFP or LV-shNogoA. In cultures transduced with LV-shNogoA, either alone or in combination with LV-Cre, NogoA protein levels were depleted by ≥90%. In the same lysates, GluA1 and S6K were significantly reduced only upon LV-shNogoA transduction, but not following genetic deletion of *Nogo* with LV-Cre (**Figure 1C-D**). Moreover, in LV-shNogoA-transduced cultures alone, we detected a significant decrease in the inhibitory pre-synaptic marker GAD-65. A robust increase in the activity-dependent transcription factor Npas4, previously shown to regulate inhibitory synaptogenesis (Lin et al., 2008), was observed only in cultures transduced with LV-shNogoA, regardless of the presence or absence of LV-Cre (**Figure 1C-D**). Thus, robust changes in synaptic proteins are observed following shRNA-mediated NogoA knockdown, but not following genetic deletion.

In a further attempt to link the observed changes in synaptic proteins to loss of Nogo signaling, we prepared hippocampal cultures from mice deficient for the Nogo receptors *Nogo receptor 1* (*Ngr1)* and *Paired immunoglobulin-like receptor b* (*Pirb)* (*Ngr1^−/−^;Pirb^−/−^*), and mice lacking all three *Ngr* family members (*Ngr1^−/−^;Ngr2^−/−^;Ngr3^−/−^*) (Atwal et al., 2008; Dickendesher et al., 2012; Fournier et al., 2001; Lee et al., 2008; Raiker et al., 2010a; Zheng et al., 2005). WB analysis for GluA1 and pS6K revealed no differences when compared to parallel-processed wildtype cultures. However, when wildtype and Nogo receptor knockout hippocampal cultures were transduced with LV-shNogoA, a significant reduction in GluA1 and pS6K was observed (**Figure S1E**). To rule out potential neurotoxic or synaptotoxic effects associated with LV-shNogoA transduction, we probed lysates for the post-synaptic density scaffolding protein PSD-95 and the glutamatergic NMDA receptor subunit GluN2B. WB analysis shows that these proteins were not altered following LV-shNogoA transduction, indicating that changes in synaptic proteins are likely not due to vector-associated toxicity (**Figure 2A-B**). Similarly, qRT-PCR studies revealed that in LV-shNogoA-transduced neurons, mRNA levels for GluA1 (*Gria1)*, GluA2 (*Gria2*), and S6K (*Rps6kb1*) are reduced, while expression for the NMDA receptor subunits GluN1 (*Grin1*) and GluN2b (*Grin2b*) or the post-synaptic scaffolding protein Homer-1 (*Homer1)* remained unchanged (**Figure 2C**).

**Figure 2.**
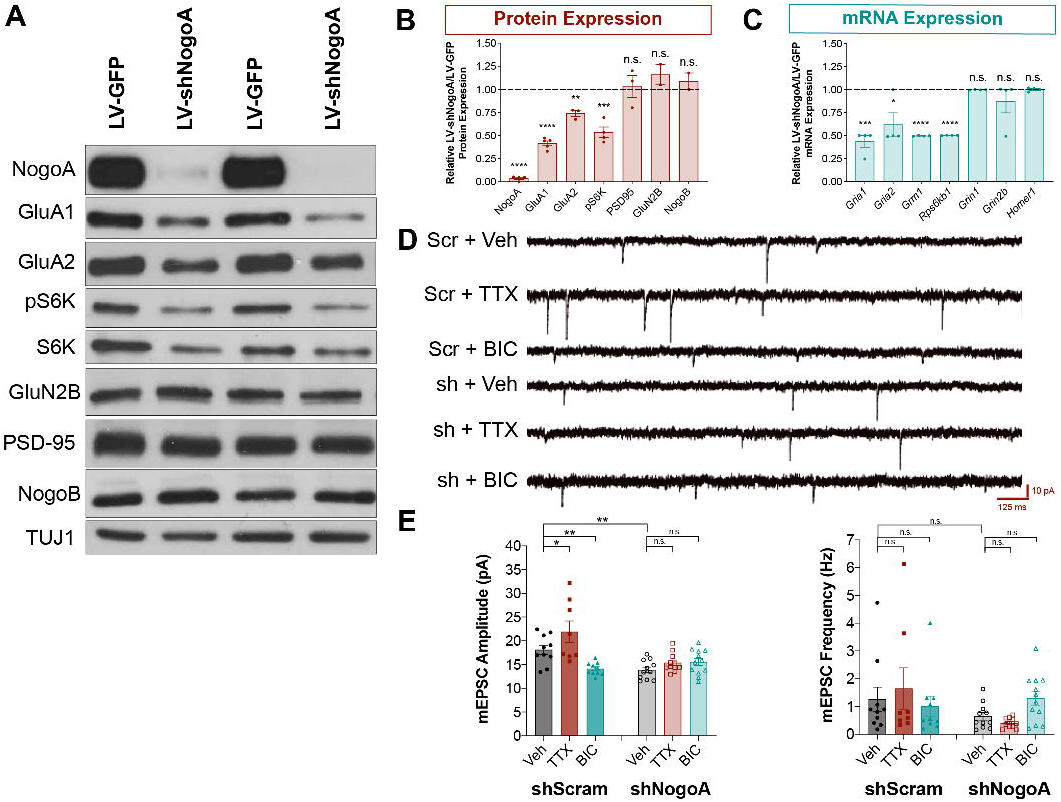
**(A)** Western blot analysis of primary rat hippocampal neurons transduced on DIV10 with LV-GFP or LV-shNogoA, and lysed on DIV17. **(B)** Quantification of protein levels, normalized first to TUJ1, then to the LV-GFP transduced cultures, *n* = 2-5 biological with 3 technical replicates each. **(C)** Quantification of qRT-PCR analysis of the select synaptic gene products as shown in (B), normalized first to *Gapdh*, then to LV-GFP. *n* = 4 biological replicates. For quantification shown in (B) and (C), \**p* < 0.05; \*\**p* < 0.01; ***p < 0.001; \*\*\**p* < 0.0001, as assessed by unpaired two-tailed Student’s *t*-test. n.s., not significant. **(D)** Representative traces of DIV14 mEPSCs recorded from neurons sparsely transfected on DIV12 with shScrambled (Scr) or shNogoA (sh) plasmids. Cultures were treated with vehicle (veh), 1μM TTX, or 10μM bicuculline (BIC) for 24hrs. Scale bars = 10pA (vertical), 125ms (horizontal). **(E)** Quantification of mEPSC amplitude (pA) and frequency (Hz) from *n* = 8-10 GFP^+^ neurons per group. All data are presented as mean ± SEM. \**p_adj_* < 0.05; \*\**p_adj_* < 0.01 as assessed by ordinary one-way ANOVA, followed by uncorrected Fisher’s LSD multiple comparisons test. n.s., not significant.

### LV-shNogoA disrupts spontaneous excitatory and inhibitory synaptic activity

Next, to determine whether biochemical changes result in altered synaptic function, we performed single cells recordings on sparsely transfected primary hippocampal neurons. On DIV12, cultures were transfected with shScram or shNogoA plasmids. For identification of transfected cells, shRNA plasmids also contained a GFP expression cassette. Depletion of NogoA in GFP^+^ neurons was verified 2 and 4 days following transfection (**Figure S1F**). On DIV14, miniature excitatory post-synaptic currents (mEPSCs) were recorded. We found that compared to shScram transfection, shNogoA leads to a significant reduction in baseline mEPSC amplitude, but not frequency (**Figure 2D-E**). Importantly, homeostatic synaptic scaling (Turrigiano et al., 1998) in response to chronic tetrodotoxin (TTX) or bicuculline (BIC) treatment is only observed in shScram-transfected neurons, but not in shRNA-transfected neurons (**Figure 2D-E**). This suggests that in primary neurons, shNogoA functions cell-autonomously to reduce baseline mEPSC amplitude and to block homeostatic scaling. Furthermore, shNogoA transduction results in a strong reduction of GAD-65 and GAD-67, suggesting dysregulation of inhibitory synapses (**Figure 3A-D**). Additional evidence for defective inhibitory synaptogenesis stems from the strong upregulation of Npas4 (**Figures 3A-B, S2C, S2E**). In contrast to global transduction with LV-shNogoA, sparsely transfected neurons did not show any change in GAD-65 or Npas4, as assessed by immunofluorescence staining of GFP^+^ shNogoA-transfected neurons (**Figure S2A, S2B, S2D**). Moreover, recordings of miniature inhibitory post-synaptic currents (mIPSCs) from LV-shNogoA-transduced neurons revealed a significant decrease in frequency, but not amplitude (**Figure 3E-F**). Taken together, these studies show that shNogoA transduction leads to biochemical changes at both excitatory and inhibitory synapses, with concurrent functional defects in synaptic transmission.

**Figure 3.**
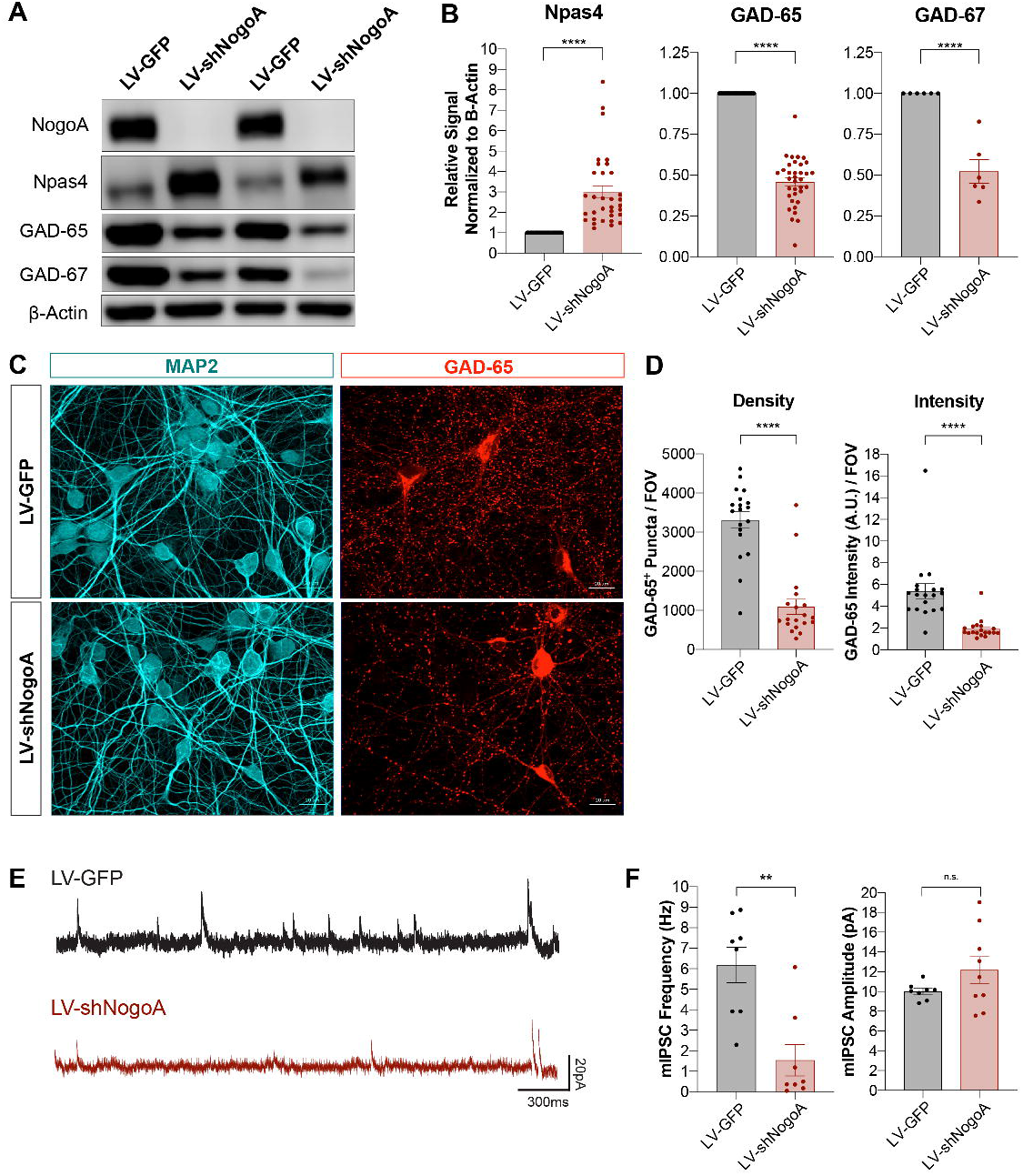
**(A)** Western blot analysis of rat primary hippocampal neurons transduced with LV-GFP or LV-shNogoA on DIV3, and harvested on DIV14. **(B)** Quantification of Npas4, GAD-65 and GAD-67 normalized first to β-Actin and then to LV-GFP. Data are presented as mean ± SEM from *n* = 33, *n* = 35, and *n* = 6 biological replicates for Npas4, GAD-65, and GAD-67, respectively. \*\*\*\**p* < 0.0001, as assessed by unpaired two-tailed Student’s *t*-test. **(C)** Immunofluorescence images of hippocampal cultures subjected to the same experimental timeline as described in (A). GAD-65 (red) marks inhibitory pre-synaptic sites and MAP2 (cyan) delineates somatodendritic neuronal morphology. Scale bar = 20μm. **(D)** Quantification of GAD-65 puncta density and intensity per field of view (FOV). Data from *n* = 19 FOV/group are shown as mean ± SEM. \*\*\*\**p* < 0.0001, as assessed by unpaired two-tailed Student’s *t*-test. **(E)** Representative traces of miniature inhibitory post-synaptic current (mIPSC) recordings from DIV14 primary hippocampal neurons globally transduced with LV-GFP or LV-shNogoA on DIV7. Scale bars = 300ms (horizontal), 20pA (vertical). **(F)** Quantification of mIPSC frequency and amplitude. Data from *n* = 8-9 neurons are presented as mean ± SEM. \**\*p* < 0.01, as assessed by unpaired two-tailed Student’s *t*-test. n.s., not significant.

### Synaptic phenotypes can be dissociated from shRNA-mediated NogoA knockdown

Alignment of the shNogoA sequence with the rat and mouse transcriptomes did not identify any likely off-target transcripts, i.e. gene products complementary to the shRNA guide strand with one or two mismatches (data not shown). To ask whether the off-target effects we observed with shNogoA are species-specific, we compared regulation of synaptic proteins in primary rat and mouse hippocampal neurons. In both rat and mouse hippocampal neurons, LV-shNogoA transduction resulted in very similar downregulation of GluA1, S6K, and GAD-65 (compare **Figures 1C-D**, 2A-B, and **3A-B**). LV-shNogoA-transduction of both rat or mouse cultures leads to a robust and calcium-dependent upregulation of the immediate-early gene (IEG) Npas4, which was similarly upregulated following KCl or BIC treatment (Lin et al., 2008) **(Figure S3)**. This shows that synaptic phenotypes observed in LV-shNogoA-transduced neurons are robust but not species-specific.

The seed region of siRNA, located between the 2^nd^ and 7^th^/8^th^ nucleotide in the 5’-region of the guide (antisense) strand, is particularly important for target mRNA recognition (Kamola et al., 2015). To determine whether synaptic phenotypes are shNogoA seed sequence-specific, we introduced either two or five point mutations in the seed region to generate an increasing number of mismatches (**Table 1**). The corresponding LV vectors, henceforth denoted as LV-shNogoA-2mt and LV-shNogoA-5mt, were generated and used for transduction of primary hippocampal neurons (**Figure 4A**). As predicted, LV-shNogoA resulted in complete, LV-shNogoA-2mt in partial, and LV-shNogoA-5mt in negligible knockdown of NogoA (**Figure 4B-D**). This shows that NogoA knockdown is seed sequence specific. Strikingly however, cultures transduced with LV-shNogoA-2mt or LV-shNogoA-5mt still exhibited a significant reduction in GluA1, S6K, GAD-65, and GAD-67, comparable to cultures transduced with the original LV-shNogoA vector (**Figure 4B-F**).Curiously, step-wise introduction of mismatches in the shNogoA seed sequence revealed novel synaptic phenotypes not observed following LV-shNogoA transduction. For example, protein levels of pre-synaptic synapsin-IIa are comparable between LV-GFP- and LV-shNogoA-transduced cultures; however, synapsin-IIa is significantly reduced following LV-shNogoA-2mt or LV-shNogoA-5mt transduction. A greater decrease in synapsin-IIa occurs as more mismatches are introduced in shNogoA sequence (**Figure 4B**).In summary, changes in synaptic protein composition in LV-shRNA-transduced cultures occur independently of NogoA knockdown; they are however, at least in part, shRNA seed sequence-dependent.

**Figure 4.**
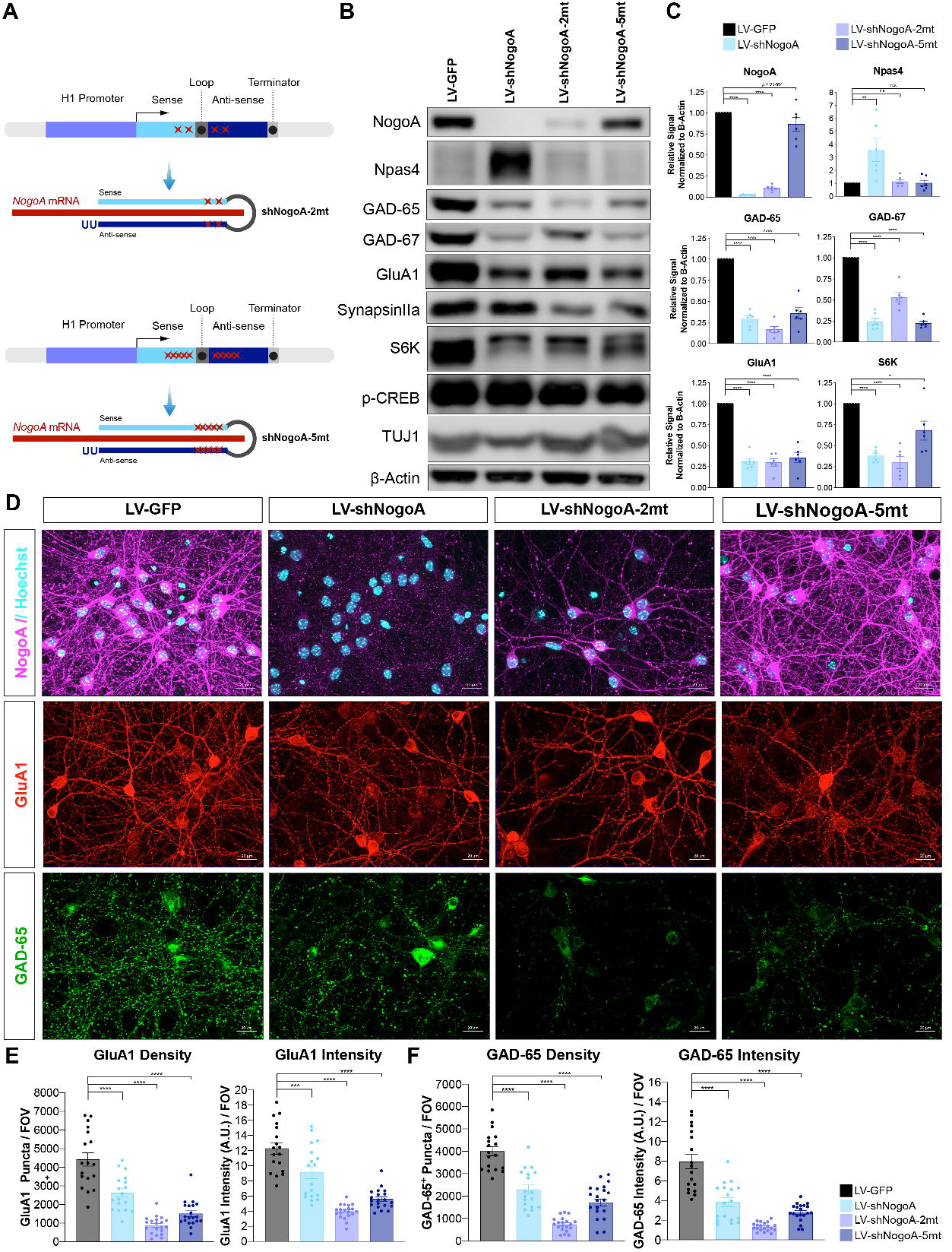
**(A)** Schematic of shNogoA seed sequence and point mutations introduced to generate two or five mismatches. **(B)** Western blot analysis of DIV14 mouse hippocampal cultures transduced on DIV4 with LV-GFP, LV-shNogoA, LVshNogoA-2mt, or LV-shNogoA-5mt. **(C)** Quantification of protein levels, normalized to β-Actin and LV-GFP. Transduction groups are labeled according to the color key. Data are presented as mean ± SEM, from *n*= 6 biological replicates. \**p_adj_* < 0.05; \*\**p_adj_* < 0.01; \*\*\*\**p_adj_* < 0.0001, as assessed by one-way ANOVA followed by Tukey’s multiple comparisons test. n.s., not significant. **(D)** Immunofluorescence images of parallel-processed DIV14 hippocampal neurons. Neurons were immunostained for NogoA (magenta) and nuclei were labeled with Hoechst (cyan). Glutamatergic post-synaptic sites are labeled with GluA1 (red). GABAergic pre-synaptic sites are marked with GAD-65 (green). Scale bar = 20μm. **(E-F)** Quantification of GluA1 (E) and GAD-65 (F) puncta density and intensity per field of view (FOV). Transduction groups are labeled according to the color key. Data from *n* = 18-20 FOV/group are presented as mean ± SEM. \*\*\*\**p_adj_* < 0.0001, as assessed by one-way ANOVA followed by Dunnett’s multiple comparisons test.

### LV-shRNA causes global dysregulation of the neuronal transcriptome

To assess transcriptional changes at a global level, we employed next-generation RNA-sequencing. For a comparative analysis, we prepared perinatal mouse forebrain cultures for DIV4 transduction with LV-GFP, LV-shNogoA, LV-shNogoA-2mt, or LV-shNogoA-5mt. Total RNA was isolated from DIV14 cultures, and split into two groups for concurrent isolation of mRNA and small RNA (**Figure 5A**). Principal component analysis (PCA) of mRNA transcriptomes revealed close clustering for cultures transduced with the same vector, demonstrating a high degree of reproducibility (**Figure 5B**). In line with biochemical findings (**Figure 4B-C**), LV-shNogoA transduction resulted in a 96% loss of the *Rtn4/NogoA* transcript, LV-shNogoA-2mt in 85%, and LV-shNogoA-5mt in 4%, when compared to LV-GFP-transduced cultures (**Figure 5E**). Transcripts for the synaptic gene products *Gria1* (GluA1) and *Gad2* (GAD-65) were downregulated in cultures with LV-shNogoA, LV-shNogoA-2mt, and LV-shNogoA-5mt (**Figure 5E**). Next, we determined the number of differentially expressed genes (DEGs) in cultures transduced with LV-shNogoA, LV-shNogoA-2mt or LV-shNogoA-5mt, when compared to LV-GFP. DEGs were identified by setting a pre-determined threshold at ≥2 fold-change and *p_adj_* ≤ 0.01 (**Figure 5D**). LV-shNogoA resulted in 807 DEGs; LV-shNogoA-2mt in 2884 DEGs; and LV-shNogoA-5mt in 2143 DEGs, when compared to LV-GFP. Moreover, we identified 3066 DEGs between LV-shNogoA vs. LV-shNogoA-2mt, 1960 DEGs between shNogoA vs. LV-shNogoA-5mt, and 3708 DEGs between LV-shNogoA-2mt vs. LV-shNogoA-5mt (**Figure 5C**). Despite these unexpectedly large changes in the neuronal transcriptomes, there is little overlap among different experimental groups. Comparison of neuronal transcriptomes revealed only 185 DEGs that are shared among cultures transduced with LV-shNogoA, LV-shNogoA-2mt, and LV-shNogoA-5mt (**Figure 5C**). Introduction of mismatches within the shRNA seed region reversed some of the original off-target effects; but generated new sets of silenced transcripts. This shows that transcriptional changes are at least in part shRNA seed sequence-dependent. Furthermore, our studies indicate that small changes in the seed sequence can cause unexpectedly large changes in the neuronal transcriptome (**Figure 5C**). To gain insights into biological processes regulated by LV-shRNA transduction, we performed STRING reactome pathway analysis (Szklarczyk et al., 2019). Analysis of DEGs between LV-GFP and LV-shNogoA-5mt transduced cultures, where NogoA expression is comparable, revealed dysregulation of gene networks involved in *transmission across chemical synapses* and *neurotransmitter receptors and postsynaptic signal transmission* (**Figure S4A**). Reactome pathway analysis of DEGs following transduction with LV-shNogoA or LV-shNogoA-2mt in comparison to LV-GFP also identified *transmission across chemical synapses* and *neurotransmitter receptors and postsynaptic signal transmission* However, STRING analysis identified additional reactome pathways that are unique for each shRNA construct (**Table 2**). Taken together, comparative analysis of neuronal transcriptomes regulated by highly similar shRNAs revealed widespread and seed sequence-dependent, but not target-specific, changes in reactome pathways.

**Figure 5.**
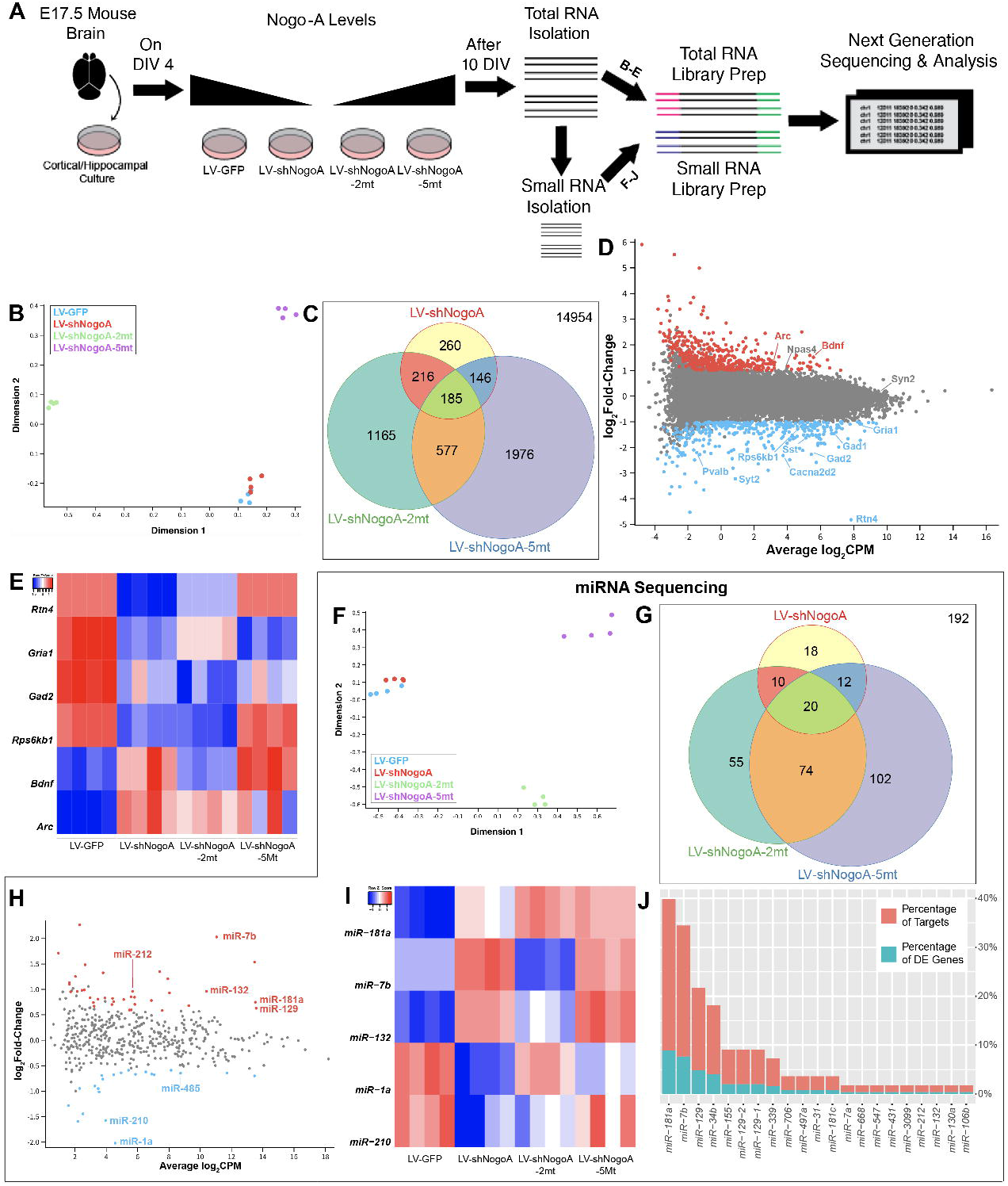
**(A)** Schematic of experimental design and timeline. On DIV4, Perinatal mouse forebrain cultures were transduced. On DIV14, total RNA was isolated, purified, and split into two groups for cDNA library preparations from total RNA (B-E) and small RNAs (F-I). Both libraries were analyzed by next generation sequencing. **(B)** Principal component analysis (PCA) of mRNA transcriptomes. Four biological replicates per LV transduction. **(C)** Venn diagram of differentially expressed genes (DEGs) with *p_ad_*_j_ ≤ 0.01 and ≥2-fold regulation compared to LV-GFP-transduced neurons. **(D)** Dot-plot shows comparison of gene expressions between LV-GFP and LV-shNogoA. DEGs with ≥2-fold regulation are shown in red (upregulated) and blue (downregulated). Several genes of interest are annotated. **(E)** Heatmap of select DEGs following indicated LV transduction. Upregulation (red) and downregulation (blue) are shown in z-scored gradient. **(F)** PCA of the parallel-processed small RNA transcriptomes. **(G)** Venn diagram of differentially expressed miRNAs with *p_adj_* ≤ 0.01 and ≥1.5-fold regulation compared to LV-GFP-transduced neurons. **(H)** Dot-plot shows comparison of miRNA expression between LV-GFP and LV-shNogoA transduced cultures. miRNAs with *p_adj_* ≤ 0.01 and ≥1.5 fold-change are marked as upregulated (red) and downregulated (blue). Some miRNAs of interest are annotated. **(I)** Heatmap of select miRNAs regulated following LV transduction. Upregulation (red) and downregulation (blue) of miRNAs are shown in z-scored gradient. **(J)** Graphs show correlation of identified miRNAs and their differentially expressed target mRNAs.

### Neuronal expression of shRNA causes dysregulation of the miRNA transcriptome

The widespread changes in gene expression were surprising, and prompted further investigations. Because large transcriptomic changes occur in cultures transduced with the original LV-shNogoA and LV-shNogoA variants with 2 or 5 mismatches in the seed region, off-target mRNA recognition and subsequent degradation seems an unlikely mechanism. Because miRNAs have been studied extensively as master regulators of post-transcriptional gene silencing in neurons (Hu and Li, 2017; Sambandan et al., 2017; Schratt, 2009; van Spronsen et al., 2013), we focused on changes in small RNA transcriptomes as a potential mechanism for the observed mRNA changes. Small RNAs of cultures transduced with LV-GFP, LV-shNogoA, LV-shNogoA-2mt, and LV-shNogoA-5mt were isolated, sequenced, and subjected to principle component analysis. Following LV-shNogoA transduction, we identified 37 miRNAs that are significantly upregulated and 22 that are significantly downregulated with a ≥1.5 fold-change and *p_adj_* ≤ 0.01, when compared to LV-GFP.

The small RNA transcriptomes between LV-GFP and LV-shNogoA are less different than those of LV-shNogaA-2mt and LV-shNogoA-5mt, which show greater divergence (**Figure 5F**). Thus, the miRNA transcriptome relationships closely mirror those observed for mRNA transcriptomes (**Figure 5B**). Between LV-shNogoA-2mt and LV-GFP, 208 miRNAs are significantly altered, and between LV-shNogoA-5mt and LV-GFP 159 miRNAs were significantly altered (**Figure 5G**). The 37 miRNAs that were upregulated following LV-shNogoA transduction were further examined to determine their target mRNAs (**Figure 5H-J**). This analysis identified miRNAs that control gene products with known synaptic functions, including *miR-181a* (Saba et al., 2012; Sambandan et al., 2017; van Spronsen et al., 2013) and *miR-132* (Remenyi et al., 2013; Scott et al., 2012). Both *miR-181a* and *miR-132* were upregulated in cultures transduced with LV-shNogoA, LV-shNogoA-2mt or LV-shNogoA-5mt, albeit to different extents (**Figure 5I**). In addition, LV-shNogoA transduction downregulated *miR-1a* and upregulated *miR-7b*, whereas transduction with LV-shNogoA-2mt or LV-shNogoA-5mt resulted in disparate changes in miRNA expression (**Figure 5I**). Strikingly, the seesaw expression pattern changes observed for these two miRNAs (**Figure 5I**) are highly reminiscent of the mRNA expression changes of *Arc* and *Bdnf*, changing in opposite directions among cultures transduced with different LV-shNogoA constructs (**Figure 5E**). These findings underscore the arbitrary nature of changes in gene expression caused by closely related shRNA constructs.

To interrogate whether there is a link between altered expressions of miRNAs and their target mRNAs, we analyzed differentially expressed mRNAs for recognition by miRNAs. Of the top 250 significantly downregulated mRNAs in LV-shNogoA-transduced cultures, 22 are known targets for *miR-181a*. In other words, 8.9% of the top DEGs are known to be regulated by *miR-181a*. This represents 30.98% of the differentially regulated mRNAs that have known miRNA targets (**Figure 5J**). Moreover, we found that the strongly but inconsistently upregulated *miR-7b* has 19 known targets, representing 7.7% of the top 250 DEGs. *miR-7b* is known to target 26.8% of the significantly regulated mRNAs that have a known miRNA in the top 250 DEGs (**Figure 5J**). *Gria1* is one of the 19 established targets of *miR-7b* (compare **Figures 5E** and **5I**). Taken together, we found that an appreciable fraction of the differentially expressed mRNAs are targets of dysregulated miRNAs.

In total, 72 of the top 250 DEGs were deemed regulated by miRNA using multimir.org version 2.2. To identify biological processes and pathways underlying known protein-protein interactions, we subjected this gene list to the STRING pathway analysis (Szklarczyk et al., 2019). The resulting interactome of miRNA-regulated mRNA products (**Figure S4B**) was less strongly correlated compared to a similar analysis including all 250 DEGs (**Figure S4A**); but was nonetheless statistically significant compared to random selection (PPI enrichment *p*-value=0.0498). This clustering analysis identified genes participating in several key biological processes and pathways, such as learning or memory (*Gria1*, *Hif1a*), modulation of chemical synapses (*Mef2c*, *Erbb4)*, and mTOR signaling pathway (*Rps6kb1*, *Map2k1*). Taken together, LV-shNogoA-driven global changes in the mRNA transcriptome mirror, at least in part, changes in the corresponding miRNA transcriptome. Moreover, some of the most prominently dysregulated miRNAs coordinate synaptic protein-protein networks.

## DISCUSSION

shRNA-mediated gene knockdown is a commonly used strategy for loss-of-function studies, as shRNAs, pre-screened for off-target mRNAs *in silico* can be developed to achieve highly efficient and long-lasting gene knockdown in neurons. However, the present study shows that neuronal shRNA expression is associated with global transcriptomic changes that are independent of the primary target. Step-wise introduction of mismatches in the seed sequence of shNogoA leads to a step-wise reduction in NogoA knockdown; yet wide-ranging changes in the neuronal transcriptome still persist. Introduction of only two mismatches (shNogoA vs. shNogoA-2mt) or three additional mismatches (shNogoA-2mt vs. shNogoA-5mt) leads to dysregulation of 3029 and 3503 genes, respectively. Surprisingly, there is little overlap in DEGs among cultures transduced with different LV-shRNA constructs. This shows that transcriptional changes are seed sequence-dependent, and not simply a reflection of LV-shRNA-associated neuronal toxicity or viral defense response triggered by double stranded RNA. Sequencing of small non-coding RNAs revealed that shRNA-elicited changes in miRNAs are seed sequence dependent. Computational analysis revealed that of the top 250 differentially expressed mRNAs, an appreciable fraction are targets of the most abundant and significantly dysregulated miRNAs. Because miRNAs regulate synaptic protein composition and transmission, these processes may be particularly vulnerable to shRNA target-independent disruption.

### Is shRNA expression toxic for primary neurons?

At the outset of this study, we tested commercially available and published shRNAs directed against NogoA (Zagrebelsky et al., 2010). Three of the four LV-shNogoA vectors tested achieved highly efficient gene knockdown in primary neurons. Loss of NogoA correlated with a strong reduction of select synaptic proteins. For example, GluA1 and GluA2, but not GluN2B and PSD-95, were significantly reduced. Similarly, proteins involved in inhibitory synapse formation and transmission were significantly reduced. We show that the reduction in synaptic proteins is not secondary to neuronal loss or synaptic toxicity, since LV-shNogoA-transduced neurons survived beyond two weeks following transduction. Moreover, WB analysis revealed levels of PSD-95 and neuron-specific β-III tubulin (TUJ1) comparable to parallel-processed cultures that were either not transduced or transduced with LV-GFP or LV-shScram. Independent evidence that neuronal health does not deteriorate upon LV-shNogoA transduction, comes from electrophysiological recordings of spontaneous synaptic activity. Recordings of mEPSCs and mIPSCs showed that both excitatory and inhibitory synapses are active and shNogoA transduction only affects either amplitude or frequency of events, respectively. Taken together, we provide multiple lines of evidence that synaptic defects observed in LV-shNogoA-transduced cultures are not simply a reflection of vector-associated toxicity.

### Should shRNA-mediated gene knockdown be used to study neuronal function?

Extensive validation of any shRNA construct used in any cell type is necessary to increase scientific rigor (Moore et al., 2010). The present work shows that careful validation of shRNA-elicited phenotypes may be particularly important in cell types that rely heavily on miRNA-mediated post-transcriptional regulation of protein networks. In developing neurons, synaptogenesis and synaptic transmission are prime examples of miRNA-regulated processes (Hu and Li, 2017; Schratt, 2009; Song, 2020). An important lesson learned from our studies is that the use of a scrambled shRNA is not a sufficient control; nor is the introduction of mismatches within the seed sequence of the shRNA directed against the gene under investigation. Neuronal transduction with LV-shNogoA-2mt and LV-NogoA-5mt yielded unpredictable patterns of mRNA and synaptic protein changes, sometimes replicating those observed with the original LV-shNogoA (e.g. GluA1, GAD-65, S6K), or introducing changes not previously observed (e.g. synapsin-IIa). While LV-shNogoA-2mt and LV-shNogoA-5mt both regulate a large number of neuronal gene products, including synaptic genes, gene regulation is inconsistent and sometimes occurs in the opposite direction. The unpredictable nature of shRNA-elicited changes in synaptic gene products makes it particularly challenging to come up with reliable strategies to distinguish between target dependent and independent effects.

A commonly used control experiment for phenotypes associated with shRNA-mediated gene knockdown is their rescue with an shRNA-resistant transgene. While considered the gold standard, controlling transgene expression levels and distribution can be challenging. For example, the choice of promoter to drive the transgene, kinetics and steady state levels of transgene expression, and potential uncontrolled effects of overexpressing a recombinant protein are some of the unknowns associated with this approach. Moreover, the untranslated regions of the endogenous mRNA may contain RNA transport motifs or harbor regions that are targeted by one or several miRNAs. Neuronal expression of an shRNA-resistant variant of *NogoA* resulted in very low levels of recombinant protein. We were unable to reach endogenous protein levels and carry out the rescue experiment. Because *Nogo^flox/flox^* mice are available, we decided to use genetic deletion as an independent means of validating shNogoA. This control experiment demonstrated that changes in synaptic proteins are not dependent on NogoA protein expression, and underscores the need for careful and independent validation of shRNA-elicited phenotypes. Because Cre recombination at the *Nogo/RTN4* locus leads to deletion of both neuronal splice variants of *Nogo*, i.e. *NogoA* and *NogoB*, it may be argued that genetic deletion of both isoforms results in a different phenotype than the knockdown of *NogoA* alone. However, subsequent transduction of LV-Cre-transduced *Nogo^flox/flox^* cultures with LV-shNogoA still elicited the same changes in synaptic protein expression, showing that they are independent of NogoB. In a parallel approach, we employed neuronal cultures prepared from Nogo receptor compound mutants. Changes in synaptic proteins in Nogo receptor-deficient cultures were not observed; however, appeared upon transduction with LV-shNogoA. Collectively, these studies provide further evidence that the observed phenotypes are not a result of impaired Nogo signaling.

In an independent study, shRNA constructs directed against *doublecortin (Dcx)* were subjected to further analysis to investigate differences in neocortical cell migration *in vivo*, following genetic deletion and shRNA knockdown of *Dcx* (Baek et al., 2014). The authors employed *Dcx*-targeting shRNAs and nine scrambled shRNAs, and found that four of the scrambled shRNAs significantly reduced neuronal migration comparable to *Dcx* shRNA (Baek et al., 2014). Similar to the present study, the observed defects were shRNA sequence-dependent, but not target-specific. Because of these seemingly unpredictable effects, the full scope of how expression of any shRNA affects neuronal gene expression can only be determined by sequencing of both the mRNA and non-coding RNA transcriptomes.

### Why does neuronal shRNA expression lead to global changes in the neuronal transcriptome?

Computational analysis shows that dysregulated miRNAs control differentially expressed mRNAs. Moreover, STRING pathway analysis of DEGs revealed protein-protein interaction networks implicated in biological processes and reactome pathways including regulation of ion transport and chemical synaptic transmission. This suggests that transcriptional changes observed in LV-shRNA-transduced cultures are at least in part a reflection of miRNA dysregulation. It has been reported that a specific miRNA may bind to the 3’-UTR of several different mRNAs, and therefore have the power to regulate multiple gene products at the same time (Lim et al., 2005). Because the miRNA copy numbers are drastically outweighed by the abundance of mRNAs (Bissels et al., 2009; Kye et al., 2007; Schwanhausser et al., 2011), changes in miRNA levels and composition could rapidly lead to coordinated changes in gene expression (Sambandan et al., 2017).

How does neuronal shRNA expression alter miRNA composition? Evidence exists that transgenic shRNAs compete for protein components necessary for proper miRNA processing in hepatocytes (Grimm et al., 2006). For example, exportin-5 and argonaute-2 play key roles in this process, where the former is involved in transporting shRNAs and pre-miRNAs from the nucleus into the cytoplasm, and the latter in miRNA biogenesis and cleavage of the bound target mRNA. Moreover, shRNA-induced saturation of the miRNA processing pathway has been observed following adeno-associated viral vector overexpression of shRNA directed against obesity-associated protein in the rat hypothalamus (van Gestel et al., 2014). Because we observed both upregulation and downregulation of individual miRNA transcripts, global transcriptomic dysregulation cannot be fully explained by simple disruption of the miRNA processing machinery.

Partial sequence complementary of the shRNA guide strand is known to cause off-target gene silencing. These effects may primarily be caused by targeting the 3’-UTR of mRNAs, and occur in a manner reminiscent of target silencing by miRNAs (Jackson et al., 2006). miRNA-like off-target effects by shRNA require only hexamer matches or partial sequence complementary to the seed region (Burchard et al., 2009), and thus, may be a fundamental property of siRNA-mediated gene silencing. Because miRNA-like off target effects are shRNA seed sequence-dependent, this mechanism may explain some of the LV-shNogoA, LV-shNogoA-2mt and shNogoA-5mt specific transcriptomic differences observed in the present study. More recent work revealed extensive communication among mRNAs, small RNAs, and RNA-binding proteins (RBPs). Crosstalk between RBPs that recognize seed sequences in miRNAs and siRNAs results in competition and causes off-target effects. This seed-to-RBP crosstalk is widespread in cancer cell lines (Suzuki et al., 2018), and likely also occurs in other cell types, including neurons.

### Dysregulation of the miRNA transcriptome in shRNA-transduced neurons causes synaptic defects

We found that some of the most significantly regulated (fold-change) and abundant (CPM) miRNAs orchestrate key biological processes, such as *modulation of chemical synaptic transmission* and *learning or memory*. Cross-reference comparison between mRNA and miRNA datasets narrowed down our focus to a subset of miRNAs, with known roles in synapse formation, maturation, and function. For example, we identified *miR-181a* as upregulated in cultures transduced with mutated shNogoAs. *miR-181a* is the most abundantly expressed miRNA at dendritic spines of hippocampal neurons, where expression of the mature form of *miR-181a* is induced as rapidly as 10 seconds following local stimulation by glutamate uncaging (Sambandan et al., 2017). Another activity-dependent miRNA we identified as upregulated by both mutated shNogoAs in our data sets is *miR-132*, which was shown to be processed into the mature form in a Dicer-dependent manner upon activation of CREB (Wayman et al., 2008). Moreover, LV-shNogoA transduction significantly downregulates *miR-485*, a previously characterized miRNA that regulates dendritic spine number and synapse formation in an activity-dependent and homeostatic fashion (Cohen et al., 2011). The list of dysregulated miRNA in our study also includes *miR-129* (Sosanya et al., 2013), *miR-132* (Remenyi et al., 2013; Scott et al., 2012), and *miR-212* (Remenyi et al., 2013). Taken together, our study identifies shRNA-mediated dysregulation of miRNAs, including miRNAs that control key aspects of synaptic formation, maintenance, and plasticity.

In summary, we report that neuronal overexpression of shRNA causes global transcriptomic changes independent of the shRNA sequence or target. In primary neurons, this leads to non-target-specific changes in synaptic protein composition and transmission. Introduction of two or three mismatches within the shRNA seed region results in wide-ranging and surprisingly different changes in the neuronal transcriptome. Lastly, analysis of the small non-coding transcriptome revealed that altered miRNA composition may contribute to target-independent synaptic changes observed in shRNA-transduced neurons.

## EXPERIMENTAL PROCEDURES

### Animals

All animal procedures were approved by the University of Michigan Institutional Committee on the Use and Care of Animals (ICUCA) and performed in compliance with the guidelines developed by the National Institutes of Health (NIH). *Nogo^flox/flox^* (Meves et al., 2018), *Nogo receptor* (*Ngr1, Ngr2, Ngr3*) compound mutants, and *Pirb* mutant mouse lines (Dickendesher et al., 2012; Lee et al., 2008; Raiker et al., 2010b) were described previously.

### Primary Neuronal Cultures

Hippocampal or forebrain neuronal cultures were prepared from perinatal mouse and rat brains. For gene transfer and pharmacological treatments, see the **Supplemental Information.**

### Immunofluorescence Labeling

Neuronal cultures prepared on glass coverslips were processed for standard immunofluorescence labeling. Quantification of signal intensity and particle density were carried out using a combination of ImageJ v1.52a (NIH Bethesda, USA) (Schneider et al., 2012) and Cell Profiler v3.1.9 (McQuin et al., 2018). For technical details, see the **Supplemental Information.**

### Western Blot Analysis

Total cell lysates were prepared from neuronal cultures in chemically reducing and denaturing conditions in the presence of protease and phosphatase inhibitors. Expression levels of select proteins were assessed using standard protocol for Western blot analysis. Protein intensity was visualized and quantified with LiCor c-Digit and Image Studio Software. For details, see the **Supplemental Information.**

### Electrophysiological Recordings

Miniature excitatory and inhibitory post-synaptic currents were recorded from sparsely transfected and LV-transduced perinatal hippocampal cultures, respectively. Electrophysiological recordings were carried out as previously described (Henry et al., 2018). For details, see the **Supplemental Information.**

### RNA Sequencing

Perinatal forebrain neuronal cultures were lysed in TRIzol. Total RNA was extracted, separated into nascent RNA, mRNA and small RNA for next-generation sequencing. For details see the **Supplemental Information.**

### Statistical Analysis

Analyses were carried out using GraphPad Prism Software (Version 8.4.2). Results obtained from a minimum of three (n=3) biological replicates from independent experiments are reported as mean value± SEM.

For comparison between two groups, unpaired two-tailed Student’s *t*-test was used. For comparison amongst multiple groups, ordinary one-way ANOVA followed by multiple comparisons tests such as Dunnett’s, Tukey’s, or uncorrected Fisher’s LSD was used. Details on statistical analyses and size of datasets are provided in the figure legends. All throughout, *p* < 0.05 was considered statistically significant.

## ACKNOWLEDGEMENTS

This work was supported by the Cellular and Molecular Biology Training Grant T32GM007315 (KTB), the Ruth Kirschstein Fellowship F31NS081852 (KTB), National Science Foundation DGE #1256260 (PMG). The National Institutes of Health, R01 MH119346, R01 NS081281 (RJG), R01 NS089896, R01 NS116008, and R21 NS104774 (SI), and the Dr. Miriam and Sheldon G. Adelson Medical Research Foundation (RJG).

## AUTHOR CONTRIBUTIONS

RK, KTB, and RJG conceptualized the study. RK, KTB, PMG, TT, AC, CGF, and XF performed experiments and collected data. RK, KTB, PMG, TT, AC, and CGF analyzed data. PMG and CJ performed biostatistical analysis. RK, KTB, PMG, MAS, SI and RJG interpreted data. RK and RJG wrote the manuscript.

