## Supplemental Information for "Small-hairpin RNAs cause target-independent microRNA dysregulation in neurons and elicit global transcriptomic changes"

Figure S1

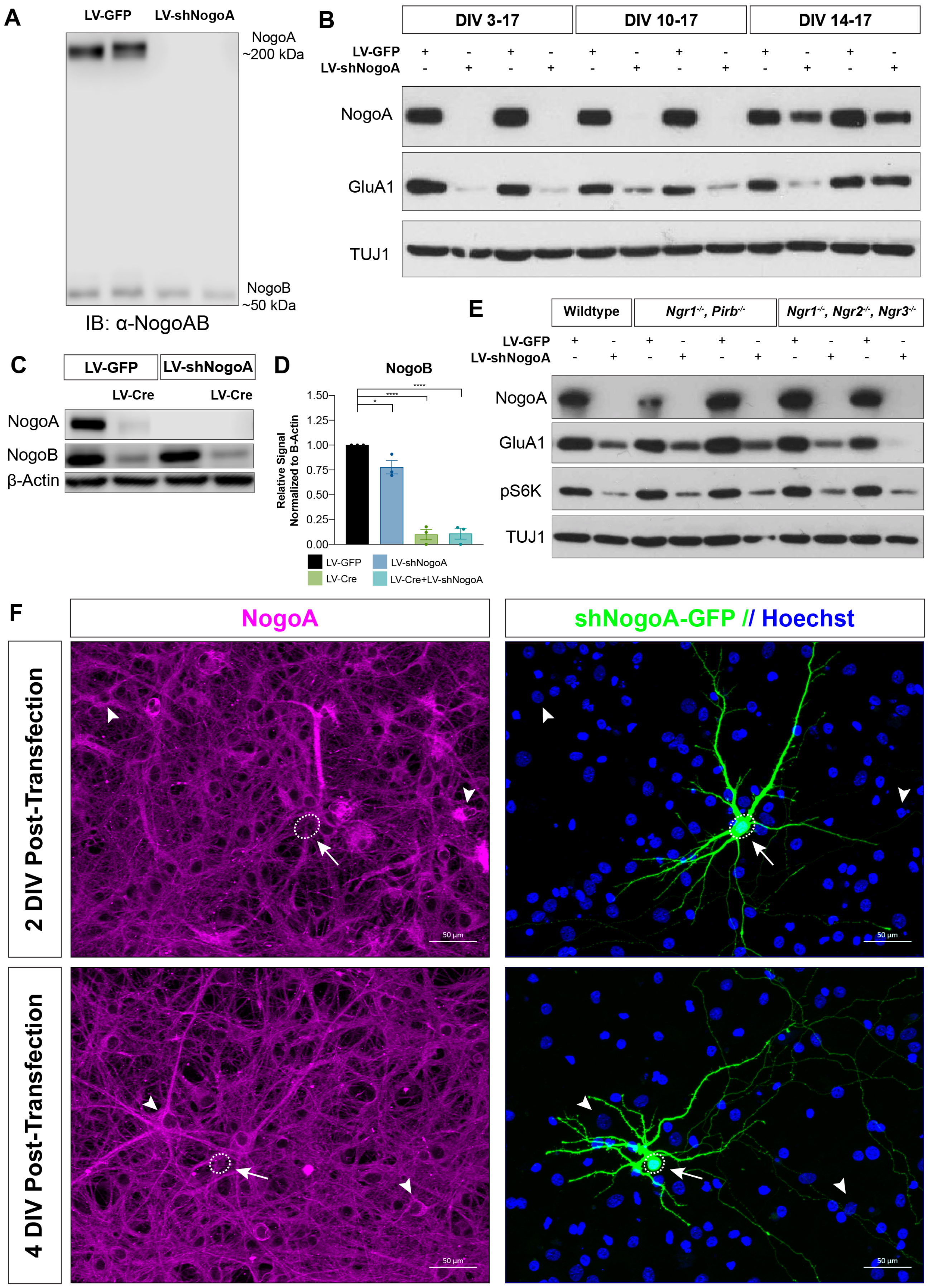
A
LV-GFP
LV-shNogoA
NogoA ~200 kDa
NogoB ~50 kDa
IB: α-NogoAB
B
DIV 3-17
DIV 10-17
DIV 14-17
LV-GFP
LV-shNogoA
NogoA
GluA1
TUJ1
C
LV-GFP
LV-shNogoA
LV-Cre
LV-Cre
NogoA
NogoB
β-Actin
D
NogoB
Relative Signal Normalized to B-Actin
1.50
1.25
1.00
0.75
0.50
0.25
0.00
LV-GFP
LV-shNogoA
LV-Cre
LV-Cre+LV-shNogoA
E
Wildtype
Ngr1-/-, Pirb-/-
Ngr1-/-, Ngr2-/-, Ngr3-/-
LV-GFP
LV-shNogoA
NogoA
GluA1
pS6K
TUJ1
F
NogoA
shNogoA-GFP // Hoechst
2 DIV Post-Transfection
4 DIV Post-Transfection
50 µm

**Figure S1. shNogoA transduction efficiently knocks down the NogoA isoform.**

**(A)** Western Blot of DIV14 lysates from E18.5 rat hippocampal cultures transduced on DIV3 with LV-GFP or LV-shNogoA, probed with the anti-NogoAB specific antibody (Bianca). NogoA (190kDa) and NogoB (55kDa) isoforms are detected in LV-GFP-transduced cultures. LV-NogoA results in selective knockdown of the NogoA isoform. **(B)** Western Blot analysis of E18.5 rat hippocampal neurons transduced with LV-GFP or LV-shNogoA on DIV3, DIV10, and DIV14. Cells were lysed on DIV17. As a loading control, the blot was probed with the neuron-specific anti-β-III tubulin antibody (TUJ1). **(C)** Western Blot analysis of *Nogo*$^{flox/flox}$ cultures transduced with LV-Cre on DIV3. On DIV6, the same cultures were transduced with LV-GFP or LV-shNogoA. **(D)** Quantification of NogoB detected by WB ($n$ = 3 biological replicates). Experimental groups are labeled according to the color key. Signal intensity was normalized first to β-actin and then to LV-GFP. Data are presented as mean ± SEM. $*p_{adj} < 0.05$; $****p_{adj} < 0.0001$, as assessed by one-way ANOVA, followed by Dunnett's multiple comparisons test. **(E)** Western Blot analysis of perinatal mouse hippocampal cultures from WT, *Ngr1*$^{-/-}$;*Pirb*$^{-/-}$ and *Ngr1*$^{-/-}$;*Ngr2*$^{-/-}$;*Ngr3*$^{-/-}$ compound mutants. Cultures were transduced with LV-GFP or LV-shNogoA on DIV10 and analyzed on DIV17. **(F)** Perinatal rat hippocampal cultures sparsely transfected with GFP-tagged shNogoA plasmid (green) on DIV12. Neurons were immunostained for NogoA (magenta). GFP$^{+}$ transfected neuronal cell bodies are delineated with a dashed circle and arrows, whereas some GFP$^{-}$ neurons are marked with arrowheads. Nuclei were stained with Hoechst (blue). Scale bar = 50μm.

Figure S2

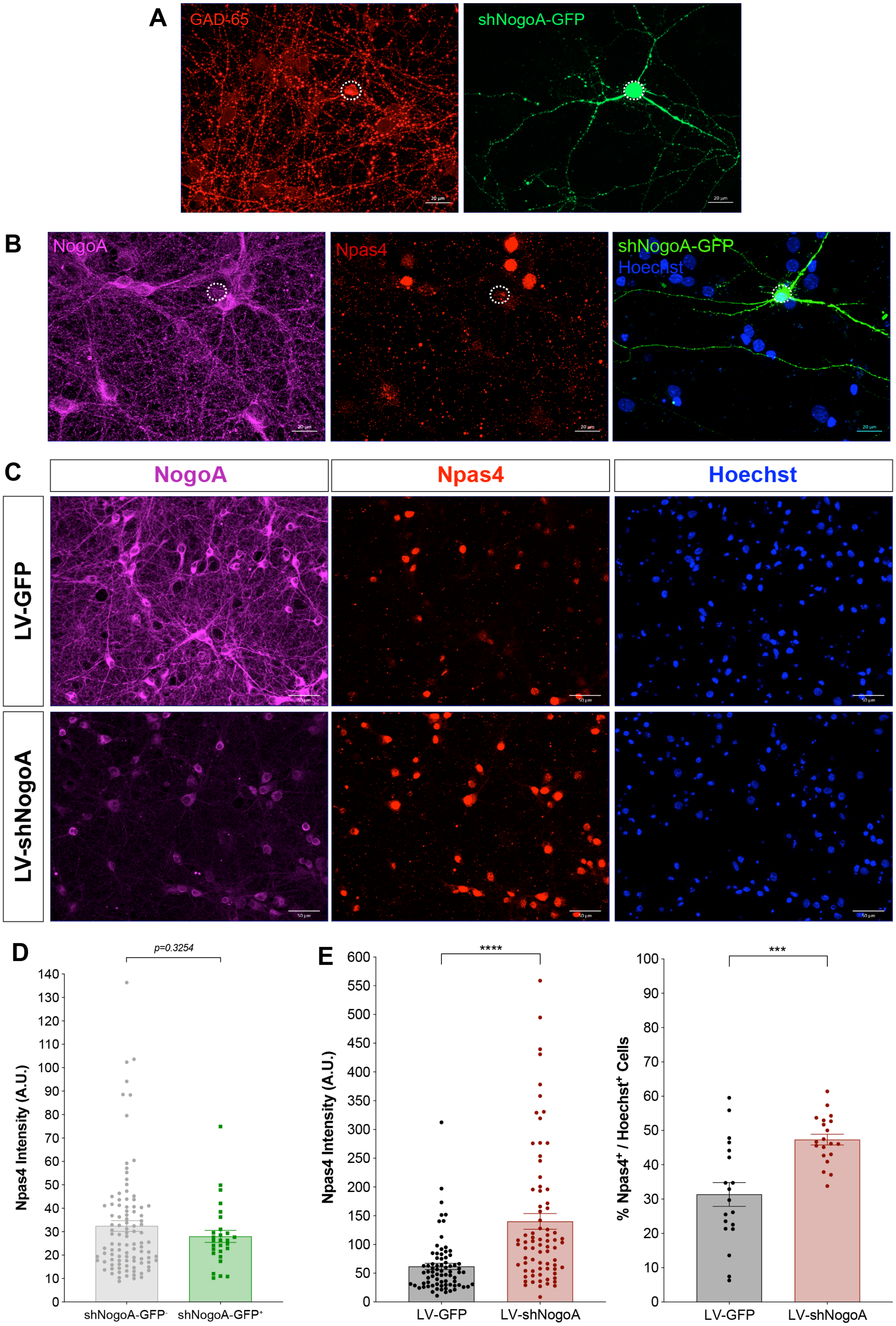

A
GAD-65
shNogoA-GFP
20 µm
B
NogoA
Npas4
shNogoA-GFP
Hoechst
20 µm
C
NogoA
Npas4
Hoechst
LV-GFP
LV-shNogoA
50 µm
D
p=0.3254
Npas4 Intensity (A.U.)
0
10
20
30
40
50
60
70
80
90
100
110
120
130
140
shNogoA-GFP-
shNogoA-GFP+
E
****
Npas4 Intensity (A.U.)
0
50
100
150
200
250
300
350
400
450
500
550
600
LV-GFP
LV-shNogoA
***
% Npas4+ / Hoechst+ Cells
0
10
20
30
40
50
60
70
80
90
100
LV-GFP
LV-shNogoA

**Figure S2. Global LV-shNogoA transduction leads to a robust upregulation of Npas4**

**(A-B)** Immunofluorescence staining of DIV14 perinatal rat hippocampal neurons sparsely transfected on DIV3 with GFP-tagged shNogoA (green, circled). Scale bar = 20μm. **(A)** GABAergic pre-synaptic sites are marked with GAD-65 (red). **(B)** Neurons were immunostained for NogoA (magenta) and Npas4 (red); nuclei were stained with Hoechst (blue). **(C)** Immunofluorescence images of DIV14 E18.5 rat hippocampal neurons globally transduced with LV-GFP or LV-shNogoA on DIV3. Neurons were immunostained for NogoA (magenta) and Npas4 (red), where nuclei were counterstained with Hoechst (blue). Scale bar = 50μm. **(D)** Quantification of nuclear (Hoechst$^{+}$) Npas4 signal intensity from GFP$^{+}$ cells following sparse transfection with shNogoA ($n$ = 28) compared to GFP$^{-}$ neighboring, NogoA-expressing cells ($n$ = 100). Deemed statistically not significant ($p$ = 0.3254), as assessed by unpaired two-tailed Student's *t*-test. **(E)** Quantifications of nuclear Npas4 expression. For percentage quantification, $n$ = 19-20 FOVs from 3 biological replicates. For intensity quantification, $n$ = 76 neurons from 3 biological replicates. Data are presented as mean ± SEM. ****$p$ < 0.0001 as assessed by unpaired two-tailed Student's *t*-test.

Figure S3

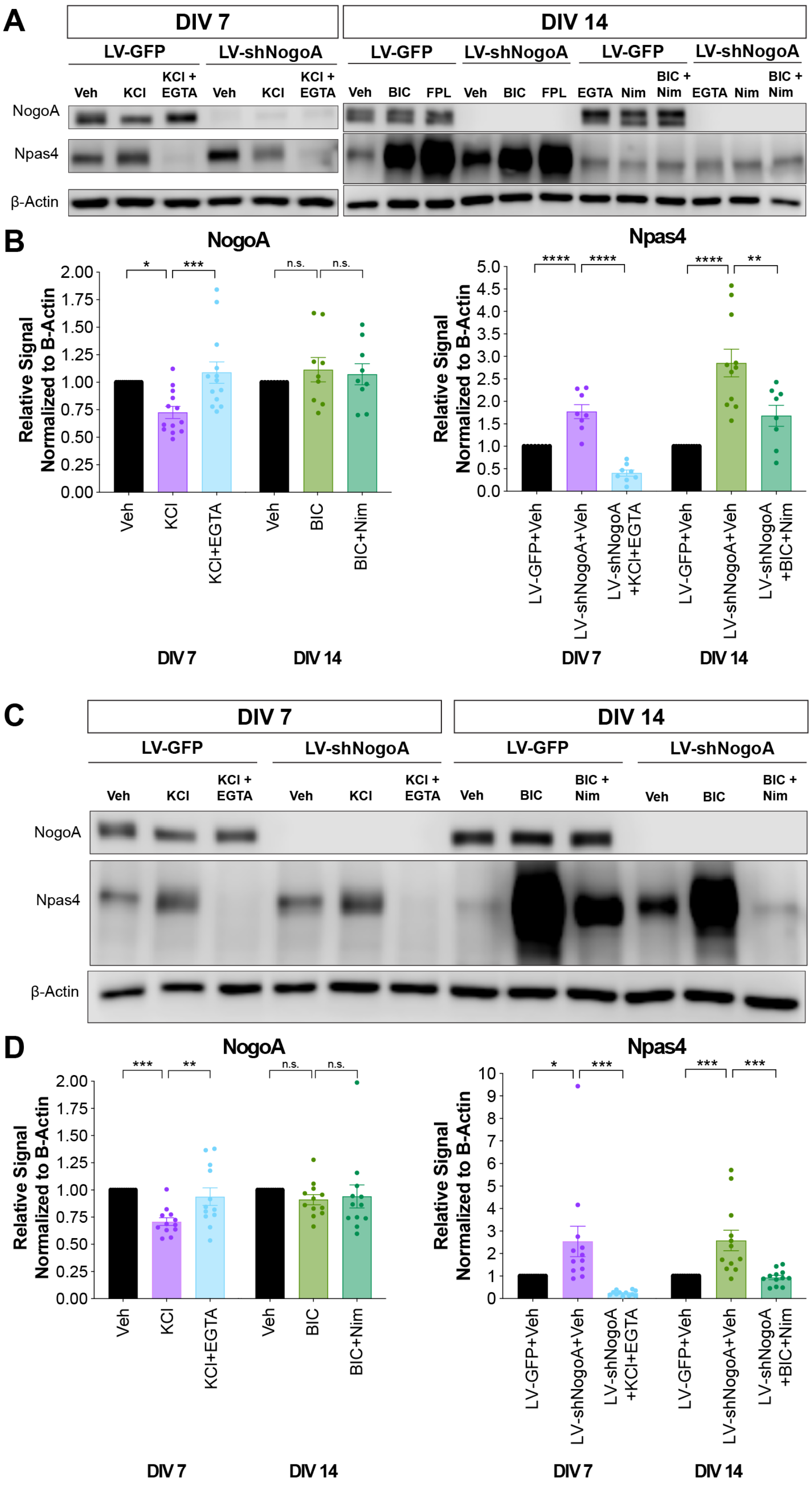

A
DIV 7
DIV 14
LV-GFP
LV-shNogoA
Veh
KCl
KCl + EGTA
BIC
FPL
EGTA
Nim
BIC + Nim
NogoA
Npas4
β-Actin
B
NogoA
Npas4
Relative Signal Normalized to B-Actin
KCl+EGTA
BIC+Nim
LV-GFP+Veh
LV-shNogoA+Veh
LV-shNogoA +KCl+EGTA
LV-shNogoA +BIC+Nim
n.s.
C
D

**Figure S3. Both rat and mouse hippocampal cultures demonstrate highly similarly phenotypes upon LV-shNogoA transduction and pharmacological modulation of neuronal activity states**

**(A, C)** Western Blot analysis of primary hippocampal neurons prepared from perinatal Sprague-Dawley rat (A) or CD1 mouse embryos (C). Cultures were transduced with LV-GFP or LV-shNogoA on DIV3, and maintained until DIV7 or DIV14. On DIV7, neurons were treated with 55mM KCl (2hrs) with or without 5mM EGTA (10mins) pre-treatment. On DIV14, neurons were treated with 50μM BIC (2hrs) or 1μM FPL64176 (2hrs) with or without 5mM EGTA (10mins) or 5μM nimodipine (1hr) pre-treatment. **(B, D)** Quantification of NogoA and Npas4 protein expression in cultures processed on DIV7 or DIV14. Data are presented as mean ± SEM from *n* = 8-13 biological replicates. *$p_{adj} < 0.05$; **$p_{adj} < 0.01$; ***$p_{adj} < 0.001$; ***$p_{adj} < 0.0001$, as assessed by ordinary one-way ANOVA followed by Dunnett's multiple comparisons test. n.s., not significant.

Figure S4

**A**

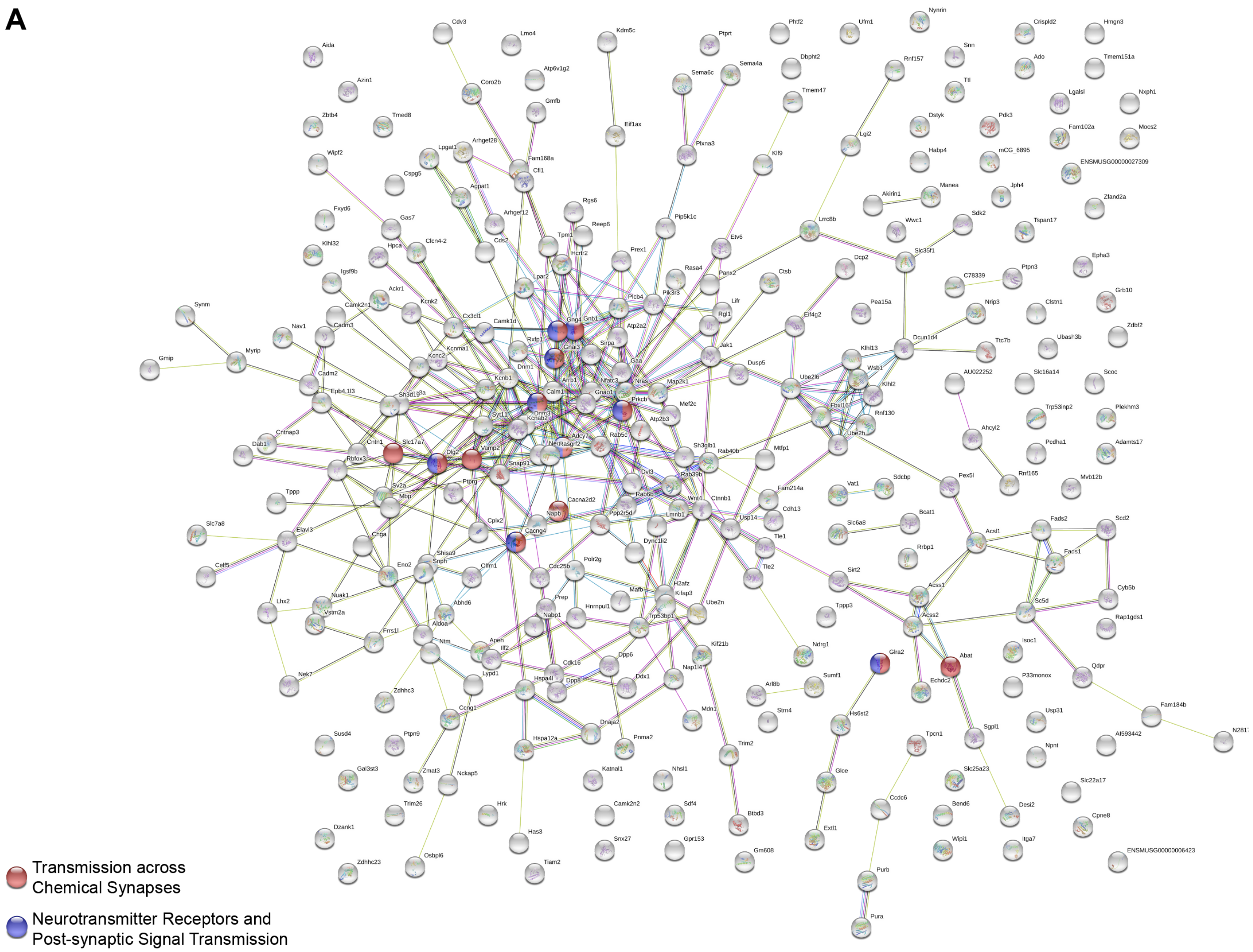
Transmission across Chemical Synapses
Neurotransmitter Receptors and Post-synaptic Signal Transmission

**B**

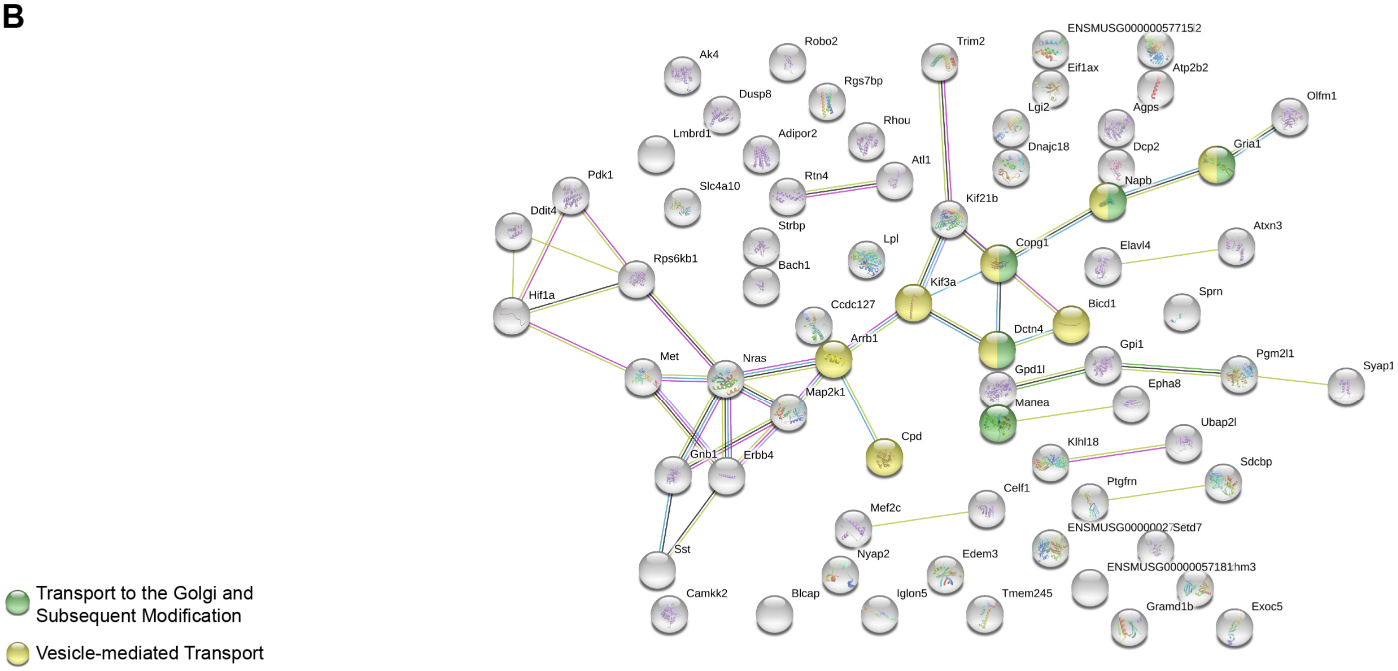
Ak4
Robo2
Trim2
ENSMUSG00000057715
Atp2b2
Eif1ax
Dusp8
Rgs7bp
Lgi2
Agps
Olfm1
Lmbrd1
Adipor2
Rhou
Dnajc18
Dcp2
Gria1
Atl1
Pdk1
Slc4a10
Rtn4
Napb
Kif21b
Ddit4
Strbp
Lpl
Copg1
Elavl4
Atxn3
Bach1
Rps6kb1
Kif3a
Hif1a
Ccdc127
Bicd1
Sprn
Arrb1
Dctn4
Gpi1
Pgm2l1
Syap1
Met
Nras
Gpd1l
Map2k1
Epha8
Manea
Cpd
Ubap2l
Klhl18
Sdcbp
Gnb1
Erbb4
Ptgfrn
Celf1
Mef2c
Setd7
Sst
Nyap2
Edem3
Camkk2
Blcap
Iglon5
Tmem245
Gramd1b
Exoc5
Transport to the Golgi and Subsequent Modification
Vesicle-mediated Transport

**Figure S4. LV-shNogoA transduction alters mRNA and miRNA transcriptomes involved in key biological processes underlying normal synapse formation and function**

**(A)** STRING analysis of the top 250 DEGs in DIV14 perinatal mouse forebrain cultures transduced on DIV4 with LV-GFP or LV-shNogoA-5mt. Lines indicate known protein-protein interactions. Proteins encoded by identified DEGs were categorized by implicated reactome pathways. **(B)** STRING analysis of the 72 DEGs with known binding by the miRNAs identified as differentially regulated following LV-shNogoA transduction. Lines indicate known protein-protein interactions. Proteins encoded by identified DEGs were categorized by implicated reactome pathways.

Table 1

| Plasmid | Source | Catalog No. | Sequence |
|---|---|---|---|
| LV-GFP | Addgene | 11795 | |
| LV-shScram | Zagrebelsky *et al*. 2010 | | GATATGAAACCCCACTTA |
| LV-shNogoA #1 | Zagrebelsky *et al*. 2010 | | AAGATTGCTTATGAAAC |
| LV-shNogoA-2mt | | | AAGATTGCTTATGACAA |
| LV-shNogoA-5mt | | | AAGATTGCTTCTAACCA |
| LV-shNogoA #2 | Open Biosystems | V2LMM_33110 | TCTCTTCCTAGTTTATGTG |
| LV-shNogoA #3 | Origene | TL711619B | CAGCAGTGTCATCCTCAGAAGGAACAATT |
| LV-shNogoA #4 | Origene | TL711619C | GATACCTTGGTAACTTATCAGCAGTGTCA |

**Table 1: Lentiviral Vector Constructs**

Table shows nomenclature, source, and sequence of all lentiviral vector (LV) constructs used in this study.

Table 2

**STRING ANALYSIS: REACTOME PATHWAYS**

**A. LV-GFP vs. LV-shNogoA**

| Pathway | Description | Count in Gene Sets | False Discovery Rate (FDR) |
|---|---|---|---|
| *MMU-112316* | Neuronal System | 18 of 300 | 0.00014 |
| *MMU-112315* | Transmission across Chemical Synapses | 13 of 187 | 0.00057 |
| *MMU-112314* | Neurotransmitter receptors and postsynaptic signal transmission | 11 of 126 | 0.00057 |
| *MMU-399721* | Glutamate binding, activation of AMPA receptors and synaptic plasticity | 5 of 25 | 0.0056 |
| *MMU-399719* | Trafficking of AMPA receptors | 5 of 25 | 0.0056 |

**B. LV-GFP vs. LV-shNogoA-2mt**

| Pathway | Description | Count in Gene Sets | False Discovery Rate (FDR) |
|---|---|---|---|
| *MMU-5576891* | Cardiac conduction | 10 of 113 | 0.002 |
| *MMU-397014* | Muscle contraction | 12 of 162 | 0.002 |
| *MMU-112316* | Neuronal System | 15 of 300 | 0.004 |
| *MMU-112315* | Transmission across Chemical Synapses | 11 of 187 | 0.0095 |
| *MMU-112314* | Neurotransmitter receptors and postsynaptic signal transmission | 9 of 126 | 0.0095 |

**C. LV-GFP vs. LV-shNogoA-5mt**

| Pathway | Description | Count in Gene Sets | False Discovery Rate (FDR) |
|---|---|---|---|
| *MMU-112316* | Neuronal System | 19 of 300 | 2.76E-05 |
| *MMU-112315* | Transmission across Chemical Synapses | 13 of 187 | 0.00071 |
| *MMU-111885* | Opioid Signalling | 8 of 64 | 0.00084 |
| *MMU-112314* | Neurotransmitter receptors and postsynaptic signal transmission | 9 of 126 | 0.0103 |
| *MMU-195721* | Signaling by WNT | 12 of 241 | 0.0127 |

**Table 2: STRING Analysis Reactome Pathways for each LV-shNogoA Transduction Group**

STRING analysis was used to identify the protein-protein interactome of differentially expressed genes (DEGs) following neuronal transduction with each of the LV-shNogoA construct variant. Table shows identified dysregulated reactome pathways, altered gene counts in the mRNA-seq data sets, and associated false discovery rates (FDRs).

**Supplemental Experimental Procedures**

***Primary Neuronal Cultures***

Time-pregnant rats and mice were euthanized with $CO_2$ followed by cervical dislocation. Embryos were extracted and kept in a sterile 10-cm petri dish (Fisher Scientific FB0875712) filled with Leibovitz's L-15 solution (Gibco 21083-027) supplemented with 1:100 Penicillin/Streptomycin (P/S) (Gibco 15140-122). Meninges were removed, and the hippocampus and neocortex (forebrain) microdissected. Tissues were rinsed in Hanks' Balanced Salt Solution (HBSS) free of $Ca^{2+}$, $Mg^{2+}$, and phenol red (Gibco 14175-095) and digested for 15-20 min at 37°C in HBSS supplemented with 0.25% Trypsin-EDTA (Gibco 15400-054) and 1 mg/mL DNAse I (Roche 10104159001). Digestion was stopped by a 5-minute incubation in ice-cold 10% fetal bovine serum (FBS) (Gibco 16000-036) in 1x Dulbecco's Modified Eagle Medium (DMEM) with high glucose and no glutamine (Gibco 11960-044). Following another wash with the same solution, tissue was washed twice in Neuronal Growth Medium (NGM): Neurobasal (Gibco 21103-049) with 25mM D(+)-glucose (Sigma G6152), 2mM Glutamax I (Gibco 35050-061), 50 units/mL penicillin and 50µg/mL streptomycin (Gibco 15140-122), and 2% B27 supplement (Gibco 17504-044). Tissue was first triturated with a single pass through a P1000 pipet tip in 1mL NGM and then triturated approximately 10x times though a fire-polished Pasteur pipette (VWR 14673-010) to fully dissociate into single cell suspension.

Trypan blue solution (Gibco 15250-061) was used for live cell counts and approximately 700,000 cells were plated per well of a 6-well plate (TPP TP92006) coated with 100µg/mL poly-D-lysine (PDL) (Sigma P7886). In parallel, 50,000 cells were seeded onto a 12-mm coverslip (Carolina Biological Supply 633029) coated with 500µg/mL poly-L-lysine (PLL) (Sigma P2636) in NGM. For electrophysiological recordings, cells were seeded onto similarly PLL-coated MatTek dishes (MatTek P35G-0-14-C) with half volume media changes every 2-3 days. For cultures that required mature synapses, cells were fed with a 1:1:1 mix of pre-warmed NGM, NCM, and astrocyte-conditioned media (ACM) supplemented with same final concentrations of B-27 and P/S, along with 2µM cytosine arabinoside (AraC) (Sigma C1768).

***Primary Astroglial Cultures***

A modified version of the previously published cortical astrocyte isolation protocol was followed (McCarthy and de Vellis, 1980). Briefly, 6 post-natal day (P)0 Sprague-Dawley rat pups were cleaned with 70% ethanol and sacrificed. Brains were removed, washed with, and dissected in Dulbecco's Phosphate Buffered Saline (DPBS) (Gibco 14287-080). Cortical tissue was microdissected and cut into small pieces and transferred into 37°C equilibrated $Ca^{2+}$- and $Mg^{2+}$-free DPBS (Gibco 14190-144) with 160U of Papain (Worthington Biochemical LS003126), 5000U of DNase I (Worthington Biochemical LS002007), and 2mg L-cysteine hydrochloride (Sigma C7477). Tissue was digested at 34°C for 45 minutes with gentle agitation. After digestion, tissue was re-suspended in 10X Low-Ovomucoid solution, containing 3g bovine serum albumin (BSA) (Sigma A8806), 3g Trypsin inhibitor (Worthington Biochemicals LS003086) prepared in DPBS. After trituration for 2 rounds of 10 passes, tissue was centrifuged before another round of trituration in High-Ovomucoid solution, containing 6g BSA and Trypsin inhibitor. After centrifugation, cells were re-suspended in

Astrocyte Growth Medium (AGM), which was a modified version of NGM, where 10% heat-inactivated donor equine serum (HyClone SH30074.03) replaced D(+)-glucose and B-27 supplement.

After filtering the cell suspension, counting viable cells, centrifuging and re-suspending the cell pellet in AGM, approximately 20 million cells were plated onto a 10μg/mL poly-D-lysine (Sigma P6407) coated 75 $cm^2$ flask (TPP TP90075) with 15mL of AGM. After a half media change with fresh AGM on DIV 1, flasks were allowed to reach confluence on about DIV3, during which time flasks were agitated vigorously until holes in the astroglial layer were created. After most contaminating non-astroglial cells were sufficiently shaken off, fresh AGM was added to the flask. On DIV5, another half-media change was carried out, with a final concentration of 2μM AraC (Sigma C1768) to inhibit mitotic fibroblasts. Before initiating the collection of ACM, astrocytes were rinsed with AGM containing 1% donor equine serum. ACM was collected every 3-5 days for approximately 1.5 months. Each collection was immediately filtered and stored at -20°C.

### *Pharmacological Treatments*

Neuronal cultures were treated with the following reagents in reported final concentrations, catalog numbers, and solvents: 55mM potassium chloride (Fisher Scientific P330) [Neurobasal], 5mM EGTA (Sigma E8145) [Neurobasal], 2μM tetrodotoxin citrate (Abcam ab120055) [dd$H_2$O], 40-50μM (+)- bicuculline (Tocris 0130) [DMSO], 200nM K252a (Calbiochem 420298) [dd$H_2$O], 10μM forskolin (Sigma F6886) [DMSO], 1μM FPL64176 (Tocris 1403) [DMSO], 5μM nimodipine (Tocris 0600) [DMSO]. For electrophysiological recordings, neuronal cultures were treated with 1μM tetrodotoxin (Tocris 1078) [dd$H_2$O] or 10μM 1(*S*),9(*R*)-(−)-Bicuculline methiodide (Sigma 14343) [dd$H_2$O]. For BrU-seq experiments, neuronal cultures were incubated with 2mM 5-Bromouridine (Sigma 850187) [dd$H_2$O].

### *Transient and Sparse Transfection*

Plasmids used for transfection are described in **Table 1**. Primary hippocampal cultures were transfected on DIV3. For each well of a 24-well plate, 0.5μg plasmid DNA, purified in endonuclease-free solutions, was mixed with 1μL Lipofectamine 2000 (Sigma L3000008) and 100μL OptiMEM (Gibco 31985070). Following a brief incubation at 37°C, this solution was mixed with NGM and used to replace culture media for 3-4 hours.

For electrophysiological studies, perinatal rat hippocampal cultures were transfected on DIV12. Neuronal cultures were first supplemented with transfection media at a 1:1 ratio (BrainBits TM500), keeping the conditioned media for after transfection. Then, 100μL transfection reagent was mixed with 1-2μL Lipofectamine 2000 (Sigma L3000008) and 100μL of transfection reagent was mixed with 1μg of plasmid DNA. Solutions were combined by gentle inversion, incubated for 20 minutes at room temperature, mixed and 200μL were added to neurons cultured on glass MatTek dishes. Cultures were returned to the incubator for 1-2 hours, and supplemented with conditioned growth media until DIV14.

***Lentiviral Transduction***

Lentiviral vectors (LV) were prepared and tittered by the University of Michigan Vector Core. For details on vector plasmids see **Table 1**. LVs were diluted in Neurobasal medium, and used at a multiplicity on infection (MOI) of 1. At a MOI 1, ~80% of primary neurons were transduced. LV transductions were carried out on DIV3-6, expect for experiments that required transduction at different developmental stages or with multiple LVs. All LV transductions were carried out in half-media changes with fresh and pre-warmed NGM.

***Immunofluorescence Labeling***

Neuronal cultures were washed with 1x Phosphate-Buffered Saline (PBS) pH 7.4, and fixed with 4% paraformaldehyde (Sigma 158127), supplemented with 4% sucrose (Fisher Scientific S5-3), for 15 minutes at room temperature. Following additional PBS washes, cells were permeabilized with 0.3% Triton X-100 (Sigma T8787) in 1x PBS, for 5 minutes. Cells were then blocked with 5% donkey serum (EMD Millipore S30), 0.1% Triton X-100 (omitted for extracellular staining) in PBS for 1 hour at room temperature, and incubated overnight at 4ºC with 1:500 dilutions of the following primary antibodies: mouse anti-NogoA (11C7, courtesy of Martin E. Schwab), rabbit anti-Npas4 (courtesy of Michael Greenberg), chicken anti-MAP2 (Abcam ab92434), mouse anti-GAD-65 (Santa Cruz sc-377145), rabbit anti-GluA1-CT (EMD Millipore AB1504).

Next day, following PBS washes, coverslips were incubated with species-specific, Cy- or Alexa fluorophore-conjugated secondary antibodies (Jackson ImmunoResearch Laboratories and Life Technologies) diluted at 1:500 in blocking buffer for 1 hour at room temperature. Nuclei were counterstained with 1:50’000 Hoechst 33342 (Invitrogen) diluted in 1x PBS. Coverslips were washed extensively and mounted in antifade Prolong Gold (Thermo Fisher Scientific P36930) on microscope slides (Fisher 12-550-15). Coverslips were then imaged with a Zeiss Apotome2 microscope equipped with an Axiocam 503 mono camera and Zen software.

For quantification of signal intensity, acquired CZI images were Apotome processed, z-stacked, and exported as high-resolution TIFF images. Images were then imported into ImageJ v1.52a (NIH Bethesda, USA) (Schneider et al., 2012), where merged channels were split into separate 8-bit images. For nuclear Npas4 signal quantification, dimensions of neuronal-like, Hoechst-labeled nuclei were outlined using the freehand drawing tool using the DAPI channel, approximating to ~3-4μm$^2$ in area. The same outlined shapes were transposed onto the dsRed channel, where Npas4 signal was acquired and stored. For each field of view, a randomly determined and placed box was used for background signal quantification. This background signal was subtracted from calculated mean value, where the resulting product was then multiplied by the nuclear area for final signal intensity value. For global GAD-65 and GluA1 signal quantification, the mean value for signal intensity from the entire field of view was measured. Signal intensities collected from multiple fields of view from each coverslip, experimental and biological replicates were pooled together for statistical analysis.

For quantification of global GAD-65 and GluA1 density, acquired CZI images were Apotome processed, z-stacked, and exported as high-resolution TIFF images. Images were then imported into ImageJ v1.52a (NIH Bethesda, USA) (Schneider et al., 2012), where merged channels were split into separate 8-bit images. Both signal intensity and size thresholds were set and applied for each field of view. Accepted particle number was measured for the entire field of view. Density values were collected from fields of view from each coverslip, experimental and biological replicates were pooled together for statistical analysis.

For quantification of percentage of Npas4-expressing nuclei, similar image processing was carried out. Single channel, 8-bit images were uploaded onto Cell Profiler v3.1.9 (McQuin et al., 2018). Using the DAPI channel, Hoechst-labeled neuronal-like nuclear area was subjected to a filter based on a singular and consistent threshold. On the dsRed channel, Npas4 signal intensity was accepted as positive signal, if and when it fell within the standard threshold range, which was determined by the mean background and signal values derived from the above-mentioned signal intensity analysis. Following streamlined analysis of multiple fields of view from each coverslip, experimental and biological replicates, percentages of Npas4-expressing, Hoechst-labeled nuclei were pooled together for statistical analysis.

***Western Blot Analysis***

Primary neuronal cultures were rinsed with PBS and lysed in Brij Lysis Buffer (10mM potassium phosphate pH 7.2, 1mM EDTA, 10mM $MgCl_2$, 0.5% NP40, and 0.1% Brij-35) or RIPA buffer (150 mM NaCl, 50 mM Tris, 1% NP-40, 3.5 mM sodium dodecyl sulfate, 12 mM sodium deoxycholate, pH 8.0), both supplemented with 50mM β-glycerophosphate (BGP), 1mM $Na_3VO_4$, and 1:100 Protease Inhibitor Cocktail (PIC) (Sigma P8340). Cells were lysed for 20-30 minutes on ice, scraped off plates, and centrifuged at 15,000rpm at 4°C for 10 minutes (Eppendorf 5430R). Supernatant was transferred to a new tube, and protein concentration was measured with a *DC* Protein Assay Kit (Bio-Rad 5000111) using a photospectrometer at 750 nm (Molecular Devices SpectraMax M5e). Supernatant was then diluted with an equal volume of 2x Laemmli sample buffer with β-mercaptoethanol (BME), boiled at 100°C for 10 minutes, and stored at -80°C. For analysis of nuclear proteins, some cultures were directly lysed in a 1:1 mixture of RIPA buffer and 2x Laemmli buffer with BME. The entirety of the lysate was boiled at 100°C for 10 minutes and stored at -80°C.

For SDS-PAGE, equal amounts of total protein (~5-10µg) for each sample were loaded per lane of a 7.5% gel. Separated proteins were transferred onto PVDF membrane (EMD Millipore IPVH00010) for 2 hours at constant 120V in cold transfer buffer: 25mM Tris, 192mM Glycine (Fisher G48), 10% methanol (Fisher Scientific A412P). Membranes were blocked in 5% blotting-grade blocker (BioRad 1706404) prepared in 1x TBS-T [TBS pH 7.4 supplemented with 0.1% Tween-20 (Sigma P1379)] for 1 hour at room temperature, and probed overnight at 4°C with the following primary antibodies diluted in 3% BSA (Fisher Scientific BP1600) in 1x TBS-T: rabbit anti-NogoAB Bianca (courtesy of Martin E. Schwab, 1:25000), goat anti-NogoAB (R&D Sytems AF3098, 1:5000), mouse anti-NogoA (courtesy of Martin E Schwab, 1:5000), mouse anti-β-actin (Sigma A5441, 1:20000), mouse anti-βIII-Tubulin (Promega G712A, 1:50000), rabbit anti-PSD-95 (EMD Millipore AB9708, 1:2000), mouse anti-SynapsinIIa (BD Pharmingen 610666, 1:5000), rabbit anti-Npas4 (courtesy of Michael E. Greenberg, 1:1000), rabbit anti-CREB p-S133 (CST 9198, 1:2000), rabbit anti-GluA1-CT (EMD Millipore AB1504, 1:500), mouse anti-GluA2 (EMD Millipore MABN71, 1:1000), rabbit anti-GluN2B (EMD Millipore 06-600, 1:1000), rabbit anti-p70 S6 Kinase (CST 9202S, 1:1000), rabbit anti-p70 S6 Kinase p-Thr389 (CST 9234L, 1:1000), mouse anti-GAD-65 (Santa Cruz sc-377145, 1:2000), mouse anti-GAD-67 (EMD Millipore MAB5406, 1:1000). PVDF membranes were then washed extensively, incubated in speciesappropriate HRP-conjugated secondary antibodies, and developed with ECL substrates. Protein band intensity was visualized and quantified in linear range with LiCor c-Digit and Image Studio Software.

***Electrophysiological Recordings***

Hippocampal neuronal cultures were prepared from P1-3 Sprague-Dawley rats, and seeded on PDL-coated MatTek dishes as previously described (Henry et al., 2018). Cultures were maintained for at least 14 DIV for synaptic maturation. Glial and cortical conditioned media were supplemented to the neuronal growth medium for the initial plating. For mEPSCs, neurons were sparsely transfected on DIV12 as described above and further treated with 1μM TTX, 20μM bicuculline, or vehicle on DIV13 for 24hrs to drive bidirectional homeostatic synaptic scaling (Turrigiano et al., 1998). For mIPSCs, neurons were globally transduced with LV-GFP or LV-shNogoA on DIV7, and all electrophysiological recordings were acquired on DIV14.

Whole-cell patch-clamp recordings were made with an Axopatch 200B amplifier for mEPSCs and with a Multiclamp 700B amplifier for mIPSCs from cultured hippocampal neurons bathed in HEPES-buffered saline (HBS) containing 119mM NaCl, 5mM KCl, 2mM $CaCl_2$, 2mM $MgCl_2$, 30mM Glucose, 10mM HEPES, pH 7.35. Whole-cell pipettes had resistance ranging 3-6 MΩ. Internal solution contained 100mM cesium gluconate, 0.2mM EGTA, 5mM $MgCl_2$, 2mM adenosine triphosphate, 0.3mM guanosine triphosphate, 40mM HEPES, pH 7.2. mEPSCs were recorded at -70mV from neurons with a pyramidal-like morphology in the presence of 1μM TTX and 10μM bicuculline, whereas mIPSCs were recorded at 0mV holding potential in the presence of 1μM TTX, 10μM CNQX, and 50μM DL-APV. All acquired traces were analyzed off-line using Synaptosoft MiniAnalysis software.

***Reverse Transcription PCR (RT-qPCR)***

Primary embryonic rat hippocampal cultures were infected on DIV9 with LVs, and maintained until DIV16. RNA was isolated using the RNeasy Mini Kit (Qiagen 74204), QiaShredder Kit (Qiagen 79654), and on-column DNase I digestion (Qiagen 79254). 1μg of RNA was used for first-strand cDNA using the SuperScript III First Strand Synthesis System (Invitrogen 18080-051). Target gene expression was assessed using $RT^2$ Profiler PCR Arrays according to the manufacturer's instructions: Rat mTOR Signaling (Qiagen PARN-098Z) and GABA & Glutamate (Qiagen PARN-152Z). Data were acquired using Applied Biosystems StepOne Plus RT-PCR Thermocycler and analyzed with StepOne Software (v.2.2.3). Cycle thresholds for each sample were normalized to *β-actin* levels. Relative mRNA expression following shNogoA transduction was compared to that of cultures with pLL3.7 control transduction. Four independent experiments were performed.

***RNA Sequencing***

For total RNA-seq, RNA was assessed for quality using the TapeStation (Agilent, Santa Clara, CA). Samples were prepared using the NEBNext Ultra II Directional RNA Library Prep Kit for Illumina (E7760L), Ribo depletion Module NEBNext rRNA Human/Mouse/Rat (E6310X), and NEBNext Multiplex Oligos for Illumina Unique dual (E6440L) (NEB, Ipswitch, MA). The rRNA-depleted RNA was then fragmented 10 minutes determined by RIN (RNA Integrity Number) of input RNA as per protocol, and copied into first strand cDNA using reverse transcriptase and dUTP mix. Samples underwent end-repair and dA-tailing step followed by ligation of NEBNext adapters. Products were purified and enriched by PCR to create the final cDNA library. Final libraries were checked for quality and quantity by TapeStation (Agilent) and qPCR using Kapa's library quantification kit for Illumina Sequencing platforms (KK4835) (Kapa Biosystems, Wilmington MA). Samples were pooled and sequenced on the Illumina NovaSeq S4 Paired-end 150bp, according to manufacturer's recommended protocols.

A total of 53-82 million reads were obtained per sample. Trim_galore 0.6.0 (github.com/FelixKrueger/TrimGalore, a wrapper around, Cutadapt version 1.15) (Martin, 2011) was used to trim the Illumina standard adapter and remove low quality bases (Phred score 20). Both reads of the pair were removed if either read had fewer than 20bp remaining. Fewer than 5% of bases were lost. STAR version 2.7.2 (Dobin et al., 2012) was used with the standard ENCODE options to align reads to the GRCm38 reference sequence; all samples had greater than 83% alignment. RSeQC (Wang et al., 2012) was used to calculate read distributions across genome features; greater than 75% of mapped reads were in an exon or 3' region. featureCounts from the Rsubread Bioconductor (Liao et al., 2019) package was used along with the comprehensive genecode.vM23.annotation GTF to summarize reads. Multimapped reads were ignored per featureCounts defaults. Counts were preprocessed using the Bioconductor package edgeR (McCarthy et al., 2012), normalized and scaled using the weighted trimmed mean (TMM) technique (Robinson and Oshlack, 2010) and further prepared for linear modeling using the Voom method from the limma package (Law et al., 2014). Linear models are then fit to Log counts per million (CPM) values using the standard limma empirical Bayes method and contrasts of interest are extracted.

For small RNA-seq, RNA was assessed for quality using the TapeStation (Agilent, Santa Clara, CA) using manufacturer's recommended protocols. Samples were prepared using the NEBNext Multiplex Small RNA Library Prep Set for Illumina (E7300L). Adapters were ligated to 100ng of total RNA, which then underwent First Strand Synthesis and PCR amplification. Products are purified and size selected by Pippin Prep according to NEB protocol recommendations. Final libraries were checked for quality and quantity by TapeStation (Agilent) and qPCR using Kapa's library quantification kit for Illumina Sequencing platforms (catalog # KK4835) (Kapa Biosystems,Wilmington MA). Samples were pooled and sequenced on the Illumina NovaSeq S4 Paired-end 150bp, according to manufacturer's recommendations.

Single end small RNA fastq files were trimmed similarly to the mRNA files expect a minimum length of 18 bases was used. Trimmed fastq files were then aligned using Bowtie (Langmead et al., 2009) through the miRDeep2 software package (Friedländer et al., 2011). miRNA's were annotated using mirbase version 22 (Griffiths-Jones et al., 2007). Counts produced by miRDeep2 were then processed through the same limma pipeline as the mRNA data to find differentially expressed miRNAs. The down-regulated mRNAs and up-regulated miRNAs were used as input into the multi-MiR R package (Betel et al., 2008; Ru et al., 2014) to map miRNAs and mRNA interactions using the multimir.org version 2.2 database. The miRmapper package was used to summarize these interactions and find miRNAs with the greatest impact (da Silveira et al., 2018). miRNAs with a large impact were defined as those that regulate the greatest number of targets. In the same sense, mRNAs regulated by the greatest number of miRNAs were also of interest.
